# Modelling homeostatic plasticity in the auditory cortex results in neural signatures of tinnitus

**DOI:** 10.1101/2022.09.12.507667

**Authors:** Hannah Schultheiβ, Isma Zulfiqar, Michelle Moerel

## Abstract

Tinnitus is a clinical condition where a sound is perceived without external sound source. Homeostatic plasticity (HSP), serving to increase neural activity as compensation for the reduced input to the auditory pathway after hearing loss, has been proposed as causal mechanism underlying tinnitus. In support, animal models of tinnitus show evidence of increased neural activity after hearing loss, including increased spontaneous and sound-driven firing rate, as well as increased neural noise throughout the auditory processing pathway. Bridging these findings to human tinnitus, however, has proven to be challenging. Here we implement hearing loss-induced HSP in a Wilson-Cowan Cortical Model of the auditory cortex to predict how homeostatic principles operating at the microscale translate to the meso- to macroscale accessible through human neuroimaging. We observed HSP-induced response changes in the model that were previously proposed as neural signatures of tinnitus. As expected, HSP increased spontaneous and sound-driven responsiveness in hearing-loss affected frequency channels of the model. We furthermore observed evidence of increased neural noise and the appearance of spatiotemporal modulations in neural activity, which we discuss in light of recent human neuroimaging findings. Our computational model makes quantitative predictions that require experimental validation, and may thereby serve as the basis of future human tinnitus studies.

**Highlights:**

- We implement homeostatic plasticity (HSP) in an auditory cortex computational model
- After HSP, model behavior shows neural signatures of tinnitus
- Increased neural noise and oscillations match human neuroimaging findings
- The proposed model can serve to design future human tinnitus studies

## 1. Introduction

Tinnitus is a clinical condition where a patient perceives a sound although no external sound source is present (Malakouti et al., 2011). Often, tinnitus is only experienced temporarily as a result of acoustic overexposure. In some cases, however, tinnitus becomes chronic, which may severely decrease the quality of a patient’s life, with known comorbidities like depression and anxiety affecting 2-3% of the population (Krog et al., 2010). In spite of decades of research, the neurobiological mechanism triggering tinnitus is still debated (Schaette and Kempter, 2006). As a result, current tinnitus treatments primarily aim at addressing the individual’s cognitive and emotional response to the tinnitus percept, and do not tackle the sound percept itself (Han et al., 2009).

Tinnitus is thought to be initiated by damage to the auditory periphery, for example due to the damage of inner hair cells. Such sensorineural hearing loss (HL) results in decreased neural activity in the auditory nerve (Kotak et al., 2005). In order to maintain homeostasis, i.e., to retain a stable mean firing rate of neurons over space and time, homeostatic plasticity (HSP) is hypothesized to occur in the deprived system (Kotak et al., 2005; Turrigiano, 1999). That is, the reduced input from the cochlea and the subsequent lower signal in the auditory nerve trigger an increase in synaptic efficacy in the central auditory system: an increased excitability and weakening of (lateral) inhibitory transmission (Eggermont and Roberts, 2012). Indeed, findings from animal electrophysiology show increased spontaneous and sound-driven responses in the cochlear nucleus (Brozoski et al., 2002; Kaltenbach and McCaslin, 1996), inferior colliculus (Auerbach et al., 2014; Mulders and Robertson, 2009), and auditory cortex (Kimura and Eggermont, 1999; Seki and Eggermont, 2003) following HL. This overall increased excitability of the auditory pathway, possibly combined with increased synchrony and changes in neuronal frequency preference (i.e., tonotopic map plasticity), is thought to lead to the perception of tinnitus (Schaette and Kempter, 2012).

While findings from animal electrophysiology provide a relative uniform picture of the changes in central auditory processing that accompany tinnitus (but see e.g., Knipper et al. (2013); Rüttiger et al. (2013)), translating these findings to human tinnitus has proven extremely challenging. Using non-invasive measurement techniques, a subset of human studies observed evidence of hyperactivity in the auditory processing pathway (Lanting et al., 2008; Leaver et al., 2011; Melcher et al., 2000). However, other studies failed to observe such effects or even observed decreased responsiveness (Gu et al., 2010; Hofmeier et al., 2018; Koops et al., 2020; Lanting et al., 2014). Tonotopic map changes were shown in earlier (Mühlnickel et al., 1998; Rauschecker, 1999) but not more recent studies (Berlot et al., 2020; Koops et al., 2020; Langers et al., 2012) of tinnitus. Electro- and magnetoencephalographic (EEG/MEG) studies reported increased delta, theta, and gamma activity with tinnitus (Adjamian, 2014; Ashton et al., 2007; Sedley et al., 2012; van der Loo et al., 2009; Weisz et al., 2005; Weisz et al., 2007), but differed in their interpretation of these findings (Sedley et al., 2012). It is unclear how findings across human neuroimaging studies relate to each other, and if the neural signatures of tinnitus observed in the human brain correspond to those observed in studies employing animal electrophysiology. The challenge in translating results across studies and species may largely stem from differences in employed research methodology and the resulting variety in spatial coverage and resolution of measurements. An explanatory gap exists between the homeostatic principles operating at the microscale (as tested through animal electrophysiology) and the meso- to macroscale information provided by human neuroimaging. For example, in the case of tonal tinnitus, processing changes relevant for the tinnitus percept may be restricted to those frequency channels that match the tinnitus sound. Such frequency-specific changes can be assessed using animal electrophysiology, but the limited spatial resolution of non-invasive measurement techniques may have obscured relevant differences between tinnitus-matched and “healthy” frequency channels. Moreover, while animal models of tinnitus are mostly employed to explore tinnitus-related changes in earlier stages of the auditory processing hierarchy, human imaging studies have largely focused on auditory and non-auditory cortical areas (but see e.g., Berlot et al. (2020); Gu et al. (2010); Melcher et al. (2000)).

Computational modeling can be used to make hypothesis-based, quantitative predictions of neural data. Thereby, computational models have the potential to bridge research findings across the diverse spatial and temporal scales dictated by the employed methodology. A wide range of computational models of tinnitus has been proposed over the past decades. In line with the data available from animal electrophysiology, earlier modeling endeavors focused mostly on subcortical processing at the microscopic scale. For example, simulation of hearing loss induced HSP through changes in excitation and inhibition in a firing rate model of the auditory nerve and cochlear nucleus showed the expected overall increased model excitability. Increased excitability was shown to lead to increased spontaneous activity, which was interpreted as representing the tinnitus percept (Gerken, 1996; Schaette and Kempter, 2006, 2008, 2009, 2012). HSP was also modeled at the level of the auditory cortex (Chrostowski et al., 2011; Dominguez et al., 2006). HSP, modeled through an increased excitability, resulted in elevated spontaneous activity and an expansion of the excitatory frequency response area of model units. In the absence of spontaneous activity, travelling waves of cortical excitation could be observed (Chrostowski et al., 2011).

Existing auditory cortical models of HSP, however, consist of spiking neurons (i.e., leaky integrate-and-fire neurons; Chrostowski et al. (2011)). While such integrate-and-fire networks provide detailed information about the short-scale temporal dynamics (i.e., processes that take less than a millisecond) of a network, they can easily become very complex, posing significant computational and interpretational challenges (Dayan and Abbott, 2001). Computational demand increases rapidly when extending a network of spiking model neurons to the meso- and macroscale accessible through human neuroimaging. Firing rate models, yielding firing rates rather than action potentials as an output, represent a simpler approach for modelling neural networks. Although less detailed than networks of interconnected spiking model neurons, firing rate models may better capture the neuronal population dynamics between inhibitory and excitatory neurons while also being more computationally efficient (Dayan and Abbott, 2001; Schaffer et al., 2013). Firing rate models therefore represent an attractive approach to modeling neuronal processes occurring at the meso- and macroscale, and may be well suited to model the neuronal processes captured in human neuroimaging data.

Here we adapted one of the most widely used firing rate models, the Wilson Cowan Cortical Model (WCCM), to simulate HSP in the auditory cortex following hearing loss. The WCCM simulates neural population dynamics by describing the interactions of excitatory and inhibitory subpopulations (Wilson and Cowan, 1972, 1973) and was recently used to model healthy auditory cortical processing (Loebel et al., 2007; May et al., 2015; Yarden and Nelken, 2017; Zulfiqar et al., 2020). We extended an existing auditory cortical model (Zulfiqar et al., 2020) by implementing HSP in two ways. First, following the existing spiking neuron model of HSP in auditory cortex (Chrostowski et al., 2011), we implemented HSP through modified cortico-cortical connectivity. Second, as the auditory part of the thalamus has been proposed as a major player in tinnitus (Llinas et al., 1999; Rauschecker et al., 2010), we also implemented HSP through additional changes in thalamo-cortical connectivity. Across ranges and severity of hearing loss, both HSP implementations resulted in neural cortical signatures that were previously associated with tinnitus, including increased spontaneous and sound-driven responsiveness restricted to the frequency range of hearing loss. Model responses furthermore showed evidence of increased neural noise, and the appearance of spatiotemporal oscillations in selected model settings. We discuss these findings in light of results from neuroimaging studies. By making quantitative predictions that require experimental validation, our model results may serve as the basis of future studies into the neurobiology underlying human tinnitus.

## 2. Materials and Methods

Figure 1 outlines our methodology to simulate HL and homeostatic mechanisms. First, we used an existing model of sound processing in the auditory cortex to represent the healthy system (*Fig. 1A*). We then introduced HL in the model (*Fig. 1B)* and modified model parameters to implement HSP in two ways: through cortico-cortical modifications only (HSP-P; *Fig. 1C*) and through additional thalamo-cortical changes (HSP-P*; *Fig. 1D)*. Spontaneous neural activity was added to the model, and differences in model behaviour under healthy, HL, and HSP conditions were then evaluated by exploring responses in the absence of input sounds, to broadband sounds, and by constructing frequency tuning curves (i.e., evaluating frequency preference and selectivity). These steps are explained in detail below.

**Figure 1.**
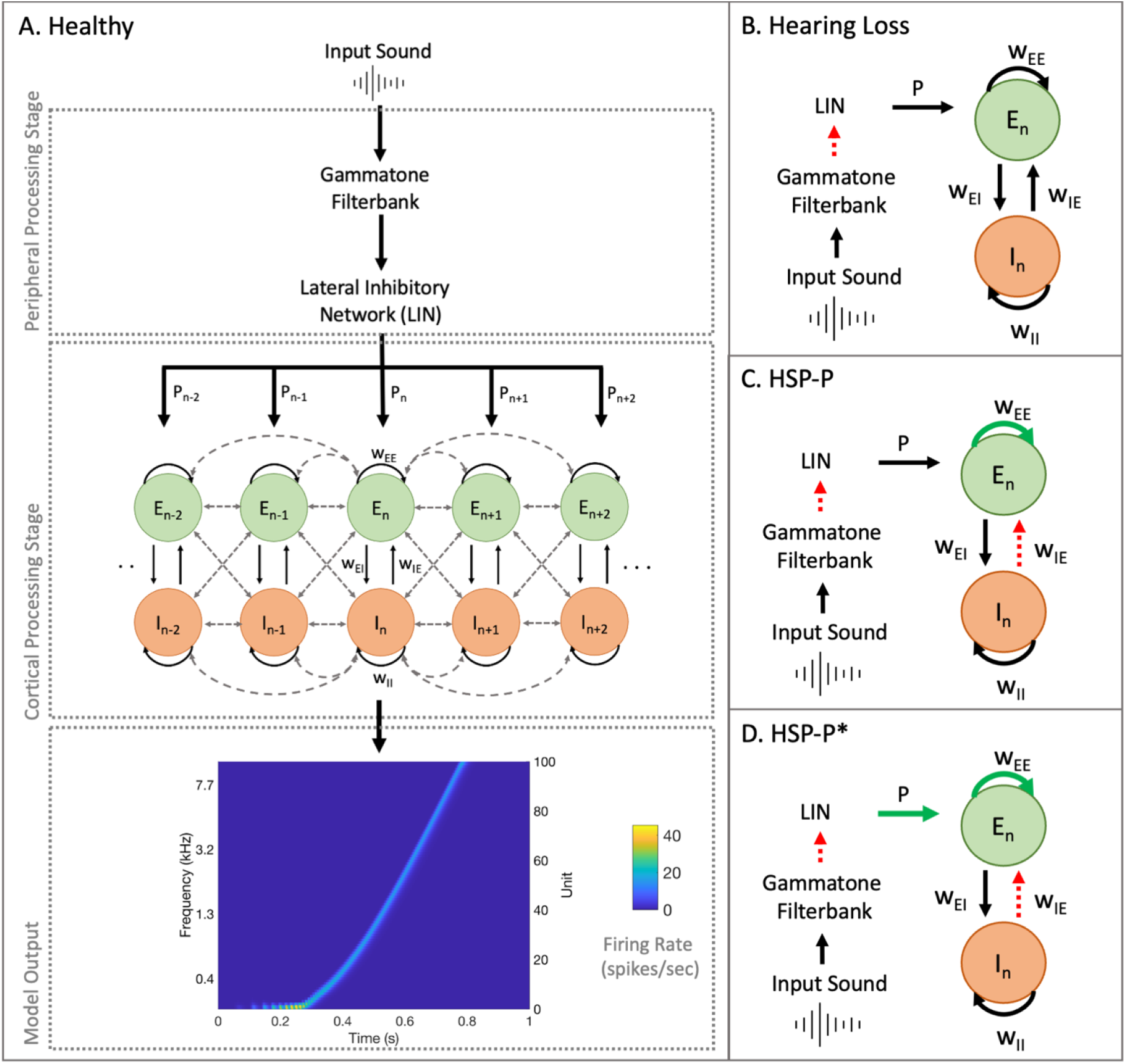
Schematic overview of the model and implemented modifications. **A**. Representation of the healthy WCCM model of the auditory cortex. In the peripheral processing stage, the input sound is first filtered through a gammatone filterbank and then spectrally sharpened using a Lateral Inhibitory Network (LIN). The output of the peripheral stage serves as an input for the cortical stage, where neuronal responses are simulated using the WCCM. Model output shows the response of the cortical processing stage to a tone sweep. **B**. HL is implemented in the model by reducing input to the LIN. **C**. Cortico-cortical HSP (HSP-P) is implemented in the model as increased excitatory-excitatory connections and decreased inhibitory-excitatory connections. **D**. Thalamo- and cortico-cortical HSP (HSP-P*) is implemented in the model as increased input from the LIN stage, as well as increased excitatory-excitatory connections and decreased inhibitory-excitatory connections. Red dashed arrows indicate a weakened connection, while green bold arrows imply strengthened connectivity.

### 2.1 WCCM of Healthy Auditory Processing

Sound processing in the auditory cortex was simulated using the model by Zulfiqar et al. (2020) (*Figure 1A*) in MATLAB (The MathWorks, Inc., v. 2020b). In this implementation, neural responses in the auditory cortex are simulated in two separate stages: a peripheral processing stage and a cortical processing stage. During the peripheral processing stage, an input sound is first passed through a Gammatone filterbank (Holdsworth et al., 1992; Ma et al., 2007; Patterson, 1986), simulating the tonotopic response of the cochlea. The filterbank consists of 100 filters with center frequencies equally spaced on an Equivalent Rectangular Bandwidth (ERB) scale, ranging from 50 to 8000 Hz. Afterwards, the sound is spectrally sharpened using a Lateral Inhibitory Network (Chi et al., 2005). To avoid boundary effects the output of the first and last unit is removed, leading to an output of 98 units (60 – 7723 Hz) of the peripheral processing stage. The cortical processing stage simulates the neuronal response in the auditory cortex using the WCCM, which comprises 98 units. Each unit is modeled by an excitatory and inhibitory unit pair, where the excitatory units receive tonotopic input from the corresponding frequency-matched peripheral stage. The interactions of the inhibitory and excitatory unit are described by Wilson (1999):

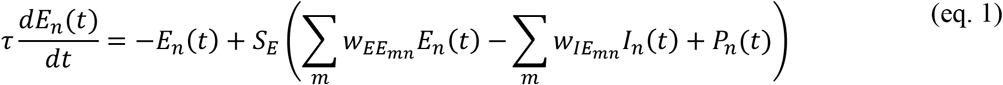

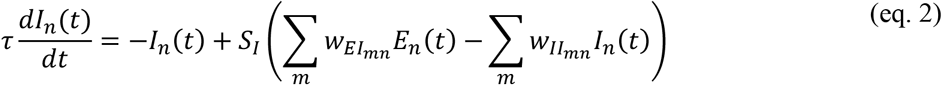

where *E*_*n*_*(t)* and *I*_*n*_*(t)* are the mean excitatory and inhibitory firing rates of neurons at tonotopic position *n* at time *t*, respectively. *P*_*n*_ represents the external input to the network and *τ* is the time constant. The four connectivity functions *w*_*ij*_ (*i*,j= E, I) represent the spatial spread of connectivity between units *m* and *n* and are described by the following decaying exponential function:

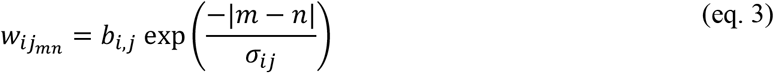

where *b*_*i,j*_ represents the maximum synaptic strength and *σ*_*ij*_ is a space constant controlling the spread of activity. Neural activity is described by the sigmoidal function *S* using a Naka-Rushton function:

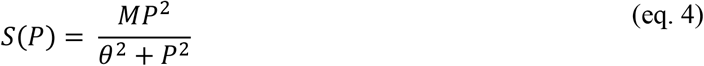

In equation (4), *M* is the maximum firing rate for high-intensity stimulus *P* and *θ* represents the semi-saturation constant, determining the point at which *S(P)* reaches half of its maximum. Values of the parameters were taken from Zulfiqar et al. (2020) and are shown in *Supplementary Table S1*.

### 2.2 Simulating Hearing Loss

To simulate HL in the model, the output from the Gammatone filterbank in response to a Gaussian white noise signal (sampling rate = 16 kHz, 1 s duration) was linearly reduced (*Fig. 1B* and *Supplementary Figure S1*). The intensity of the sound stimulus was fixed to drive the healthy model to a mean population firing rate (i.e., of all units) of 29.65 spikes/s (referred to as *E*_*n*_^*Target*^). To study the influence of HL severity on model output, different percentages of signal intensity decrease were investigated. The filterbank output was reduced by 40%, 60% and 80%, which resulted in an average firing rate of 21.77, 17.94, and 12.90 spikes/s of units within the HL region, respectively.

Additionally, HL was implemented across multiple ranges of units, specifically spanning 10, 20 or 40 model units. While HL is usually associated with higher frequencies (starting at ∼2 kHz), we implemented it in more central model units, with a BF lower than those typically affected by HL, in order to avoid model boundary effects (Pan et al., 2009). As the model parameters do not change over the tonotopic axis, similar results were expected regardless of the exact HL frequencies.

### 2.3 Simulating HSP

For an excitatory unit at tonotopic location *n*, HSP was implemented to occur when neural activity diverged from target rate (*E*_*n*_^*Target*^) in order to restore the average firing rate back to that observed in the healthy model response to broadband noise. To achieve the restoration of firing rate, two different approaches were taken.

#### 2.3.1 Updating Cortico-Cortical Connectivity (HSP-P)

The first approach only simulated modified cortico-cortical connectivity after HSP (referred to as HSP-P; *Fig. 1C*). For this, the equations proposed by Chrostowski et al. (2011) were used as a starting point and modified to avoid the violation of model constraints in WCCM (see *Supplementary Equations S1-S3* for the equations by Chrostowski et al. (2011)). The increase in synaptic efficacy of excitatory cortico-cortical synapses was achieved by increasing the value of the maximum synaptic strength 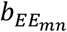in a stepwise fashion, as shown in equation (5):

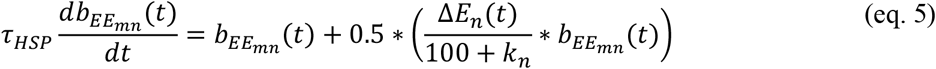

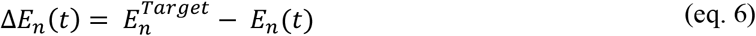

where Δ*E*_*n*_ (*t*) is the instantaneous difference from the target firing rate as is given by equation (6) and *k*_*n*_ is a scaling constant updated in steps of 50 (0,50,100, …) to allow for a slow convergence towards the target firing rate while checking at every step that model constraints are not violated. *τ*_*HSP*_ was set such that updating of the synaptic weights continued until the mean 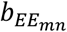value, rounded to four decimal places, did not change for five consecutive iterations.

Additionally, reduced synaptic efficacy of inhibitory cortico-cortical synapses after HSP was implemented by decreasing the value of the maximum synaptic strength 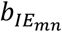in a stepwise manner:

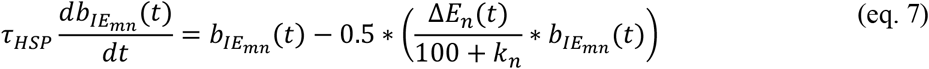

#### 2.3.2 Updating Thalamo-Cortical Connectivity (HSP-P*)

In addition to updating cortico-cortical connectivity, the second approach to HSP implementation also updated the thalamocortical input *P* (referred to as HSP-P*; *Fig. 1D*) using equations (8) and (9):

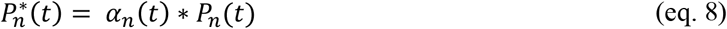

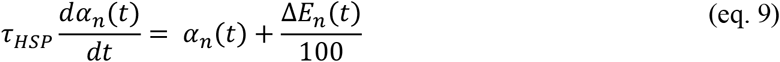

where updated input 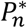is instantaneously scaled by a factor *α*_*n*_ driven by Δ*E*_*n*_ (*t*).

#### 2.3.3 Spontaneous Activity

The initial model implementation only produced output in response to an input stimulus. However, tinnitus is also perceived in the absence of an external sound, possibly related to increased spontaneous activity of neurons in the thalamus and auditory cortex. To test if the model mimicked this behavior, spontaneous activity was implemented by adding random input to the LIN output. The intensity of the random input was set to yield a mean firing rate of 2.54 spikes/s in the healthy model in the absence of an external sound (corresponding to the spontaneous firing rate in the cat auditory cortex; Seki and Eggermont (2003)). To assess whether HSP resulted in elevated spontaneous activity compared to the HL and healthy model, spontaneous firing rate (i.e., without external sound input) was compared across models.

### 2.4 Model Stability

Updating the synaptic weights of excitatory-excitatory and inhibitory-excitatory connections as part of the HSP implementations distorted the original synaptic distribution of the healthy model. Divergence of the values of 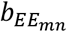from 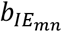eventually resulted in hyperactivity (i.e., firing rate >90 spikes/s, where the affixed maximum firing rate was 100 spikes/s). To maintain model stability (i.e., avoid hyperactivity), a two-dimensional solution space was explored, which defined the region of stability for linearly interdependent values of 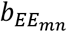and 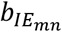(*Figure 2*). Updating input *P* (eq. 8) for the HSP-P* implementation also affected the stability constraints for a given combination of synaptic weights 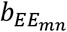and 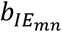. Therefore, a three-dimensional solution space was generated for HSP-P*, yielding the linearly interdependent 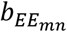and 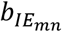values that did not cause hyperactivity in the model for varying *α*_*n*_ values used to update input *P*.

**Figure 2.**
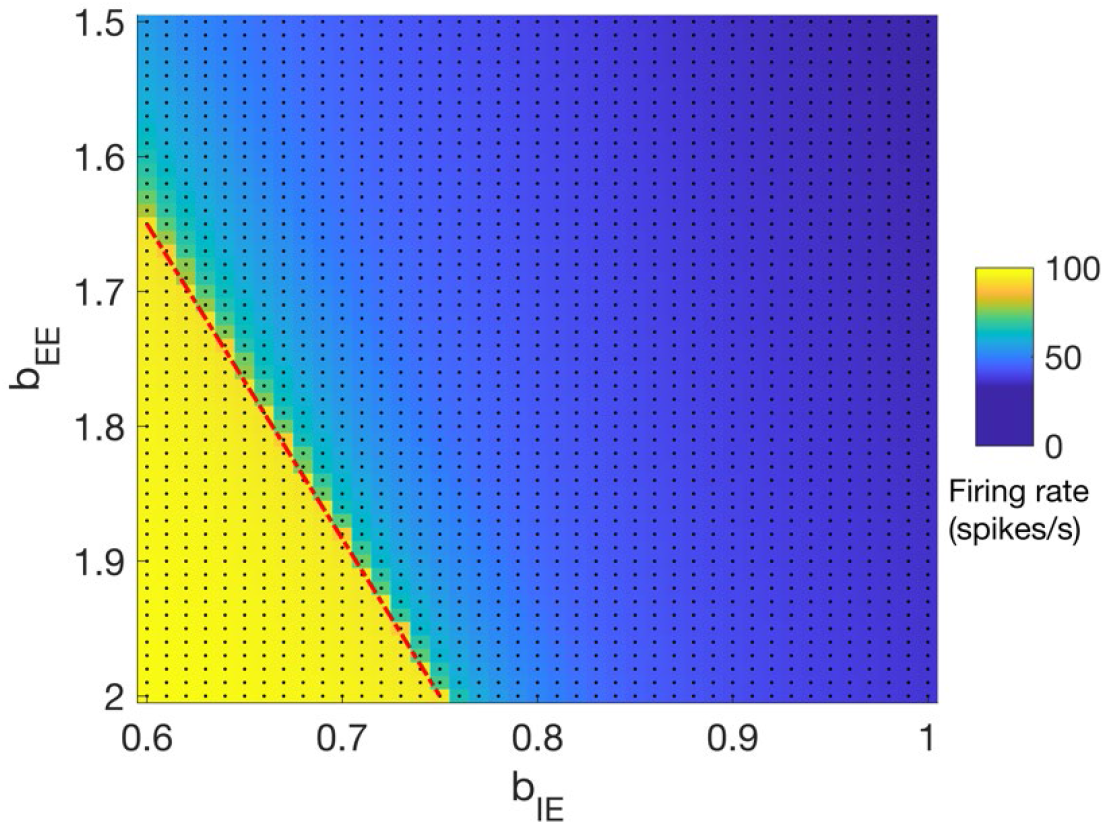
Two-dimensional solution space of 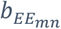and 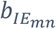values. Colors indicate the firing rate (spikes/s). The red dashed line defines the linear relationship between 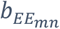and 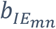values below which hyperactivity in the model was elicited (i.e., firing rate > 90 spikes/s). Black dots indicate the parameter combinations that were used for the grid search.

### 2.5 Model Evaluation

To investigate whether HSP was able to restore the target firing rate across HL ranges and severities, and to explore possible differences between the two implemented HSP approaches, model responses to Gaussian white noise as an input sound were explored. Next, to evaluate how HSP-driven changes in the model parameters affected the frequency preference and selectivity of model units, frequency tuning curves (FTCs) were constructed for all units for the healthy, HL, and HSP models. FTCs represent a unit’s response to pure tones across a range of frequencies, and thereby show to which frequency a unit responds best and how frequency-selective the unit is (i.e., the unit’s tuning width). Frequency selectivity was quantified as the bandwidth at full-width half-max (or tuning width [TW]) in octaves (Schreiner et al., 2000). To construct the FTCs, the responses of each unit to 120 pure tones (0.5 s duration) centered around the unit’s best frequency (BF) were recorded. Pure tones ranged from 0.5 octaves above until 0.5 octaves below the BF, and sampling was denser close to the BF (175 samples/octave up to ± 0.2 octaves from the BF; 83 samples/octave for the range 0.2 - 0.5 octaves from the BF). To assess how HSP affected the frequency tuning outside, inside the sloping and within the HL region, average FTCs were calculated by aligning the response of each unit around its preferred frequency, converting frequency into octaves from BF and averaging the responses of units per region (i.e., outside, sloping or within HL region). Units for which the full TW could not be computed as parts of the FTC were outside the model’s frequency response range, were excluded in the analysis.

## 3. Results

### 3.1 Model Response Restoration by HSP

First, it was evaluated whether HSP was able to restore the firing rate after HL, either through updating only cortico-cortical connectivity (i.e., HSP-P) or by additionally modifying thalamocortical input *P* (i.e., HSP-P*). To this end, the model’s responses to broadband noise were explored (*Figure 3*). The firing rate could be restored across the two implemented HSP mechanisms with 40% and 60% HL (i.e., regardless of whether thalamocortical input *P* was updated; *Fig. 3A-B*). In contrast, HSP was not able to restore the firing rate within the HL region in the case of 80% HL. While updating thalamocortical input *P* (HSP-P*) brought the resulting firing rate closer to the healthy model compared to when *P* was not updated (HSP-P; *Fig. 3C*), neither of the two implemented HSP mechanisms could fully restore the firing rate. Results were similar across HL ranges (see *Supplementary Fig. S2*).

**Figure 3.**
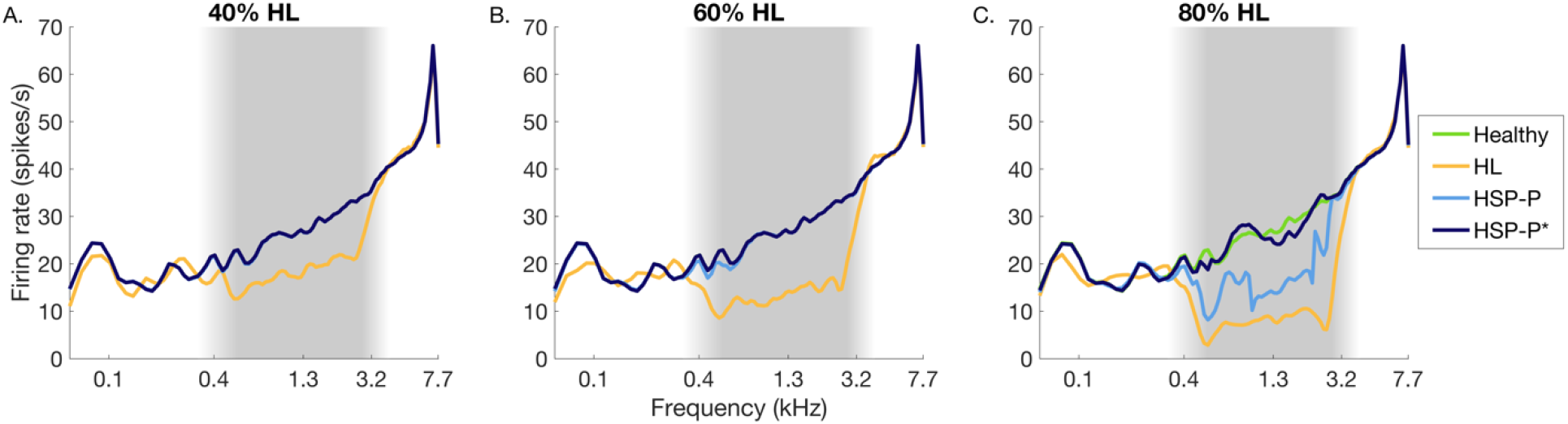
Model responses to broadband noise. Sound-evoked response in the healthy, HL, and HSP without (HSP-P) and with updated thalamocortical input (HSP-P*) are shown in green, yellow, light and dark blue, respectively. A, B and C show the responses for models with 40%, 60% and 80% HL, respectively. For all models, HL affected 40 units. Shaded grey regions delineate the HL region.

The modifications of the cortico-cortical synaptic weights (i.e., 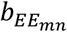and 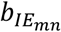) and the multiplication factor for thalamocortical connectivity *P* (*α*_*n*_)with HSP are displayed in *Figure 4*. As results were similar across HL ranges, only the results for a 40-unit HL range are shown (see *Supplementary Fig. S3-4* for the other HL ranges). As expected, across HL severity, HSP increased the maximum excitatory synaptic weight 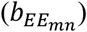, decreased the inhibitory synaptic weight 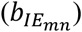, and increased the thalamocortical connectivity (through multiplication factor *α*_*n*_). Changes in model parameters with HSP were located within the HL region, and were larger when HSP was restricted to changes in cortico-cortical connectivity (i.e., HSP-P) compared to when thalamocortical input was additionally modified (i.e., HSP-P*). This pattern was expected, and confirmed that updating thalamocortical connectivity partly compensated for the response decrease after hearing loss.

**Figure 4.**
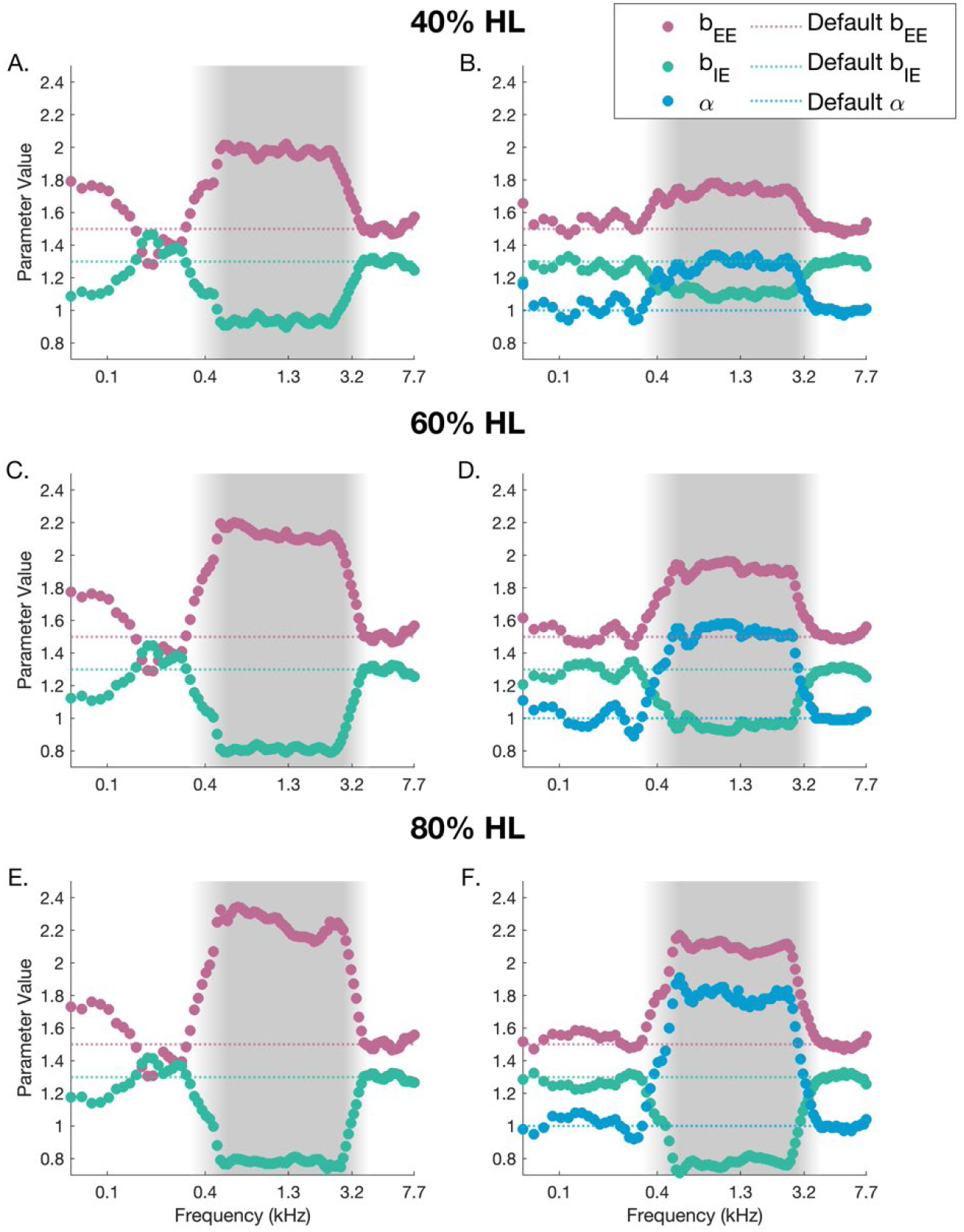
HSP-induced changes in cortico-cortical synaptic weights and thalamocortical input strength across model units. Changes in cortico-cortical synaptic weights 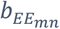and 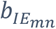are shown in pink and turquoise, respectively. Changes in the multiplication factor α_n_ for thalamocortical input P are shown in light blue. The dotted lines show the default 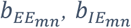, and α_n_ values of the healthy model. A, C, and E show the results for models where thalamocortical input P was not updated (HSP-P), while B, D and F display the values for models where P was updated (HSP-P*). For all models, HL affected 40 units, outlined by the shaded grey region.

Inspection of the HSP-induced changes in model parameters explained why hearing loss could not be fully restored in a subset of the scenarios. When limiting HSP to changes in cortico-cortical connectivity, in the 80% HL scenario the boundaries of 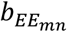and 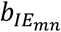values within which model stability could be ensured were reached in most of the units within the HL region, explaining why the firing rate could not be restored. In the 60% HL scenario, these boundaries were only reached for some of the units, while they were never reached for the 40% HL model. When updating thalamocortical input *P* (HSP-P*; *Fig. 4B, D, and F*), the boundaries of 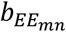and 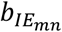 values within which model stability could be ensured were never reached. Outside the HL region, HSP did not affect model parameters. An exception to this pattern was seen for HSP-P, where 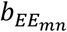and 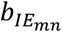values converged in units below the HL region (*Fig. 4A, C, and E*).

### 3.2 HSP Increases Spontaneous Activity and Elicits Spatiotemporal Oscillations

Next, we added spontaneous activity to the model and evaluated model responses after HSP in the absence of an input sound. In line with animal electrophysiology (Seki and Eggermont, 2003) and previous computational models of HSP (Chrostowski et al., 2011; Schaette and Kempter, 2006), we observed increased spontaneous activity after HSP (Figure 5 shows results for a HL range equal to 40 units; see *Supplementary Figures S5* and *S6* for the other HL ranges). Across HL ranges and severities, HSP caused sharp increases in spontaneous firing rate within the HL region. The spontaneous firing rate increase was clustered along the frequency axis in peaks occurring every 10-12 units. This spatially periodic activity pattern (i.e., an oscillation across the model units) is indicative of a Turing instability (Ermentrout and Cowan, 1979). The amplitude of the spontaneous firing rate peaks increased with increasing HL severity, but this increase was greater for HSP with updated thalamocortical input (HSP-P*) than for HSP where only cortico-cortical input was modified (HSP-P). As a result, while for 40% HL HSP-P resulted in the largest increase in spontaneous activity (Fig. 5A), for 80% HL the largest increase in spontaneous activity was seen for HSP-P* (Fig. 5C).

**Figure 5.**
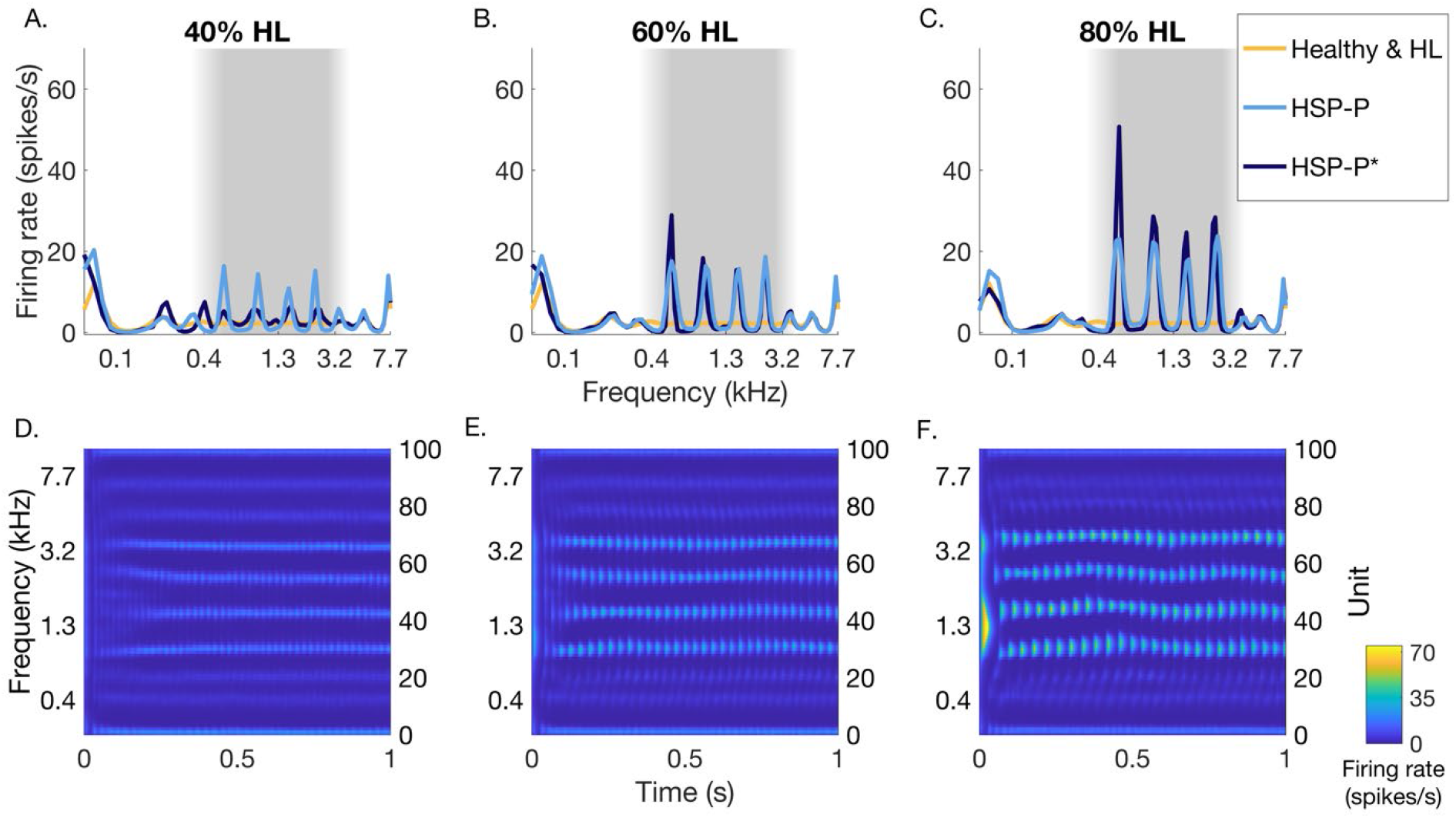
Spontaneous model activity after HSP in the absence of an input sound. A-C show the model responses averaged across time for 40%, 60%, and 80% HL, respectively. Yellow: healthy and HL responses; note that in absence of an input sound these conditions are equal, as spontaneous activity was added after the LIN stage (and was thus not affected by HL) light blue: HSP-P response without updated thalamocortical input; dark blue: HSP-P* response with updated input. D-F show the HSP-P model response over time for 40%, 60%, and 80% HL, with an average temporal oscillation rate of 45 Hz, 35 Hz, and 27 Hz, respectively. For all models, HL affected 40 units. Shaded grey regions delineate the HL region.

After a brief increase in spontaneous activity of all units within the HL region, the spontaneous activity in the model after HSP oscillated over time as well as across space (*Figure 5D-F*) indicative of a Hopf instability (Negahbani et al., 2015). The temporal oscillation rate varied between 27 Hz and 45 Hz and decreased with increasing HL severity.

### 3.3 HSP Increases Sound-Driven Activity

In order to evaluate changes in sound-driven activity after HSP, we next presented broadband white noise input to the model and evaluated responses (*Figure 6* shows results for a HL range equal to 40 units; see *Supplementary Figure S7* for the other HL ranges). Across the majority of HL levels and ranges, and across HSP implementations, HSP overcompensated the HL-driven decrease in firing rate (Table 1). That is, mean activity after HSP was elevated compared to the healthy model. This overcompensation was limited to the HL range (Figure 6), stronger when only updating cortico-cortical connectivity (i.e., stronger for HSP-P than HSP-P*), and it increased with HL range.

**Table 1.**
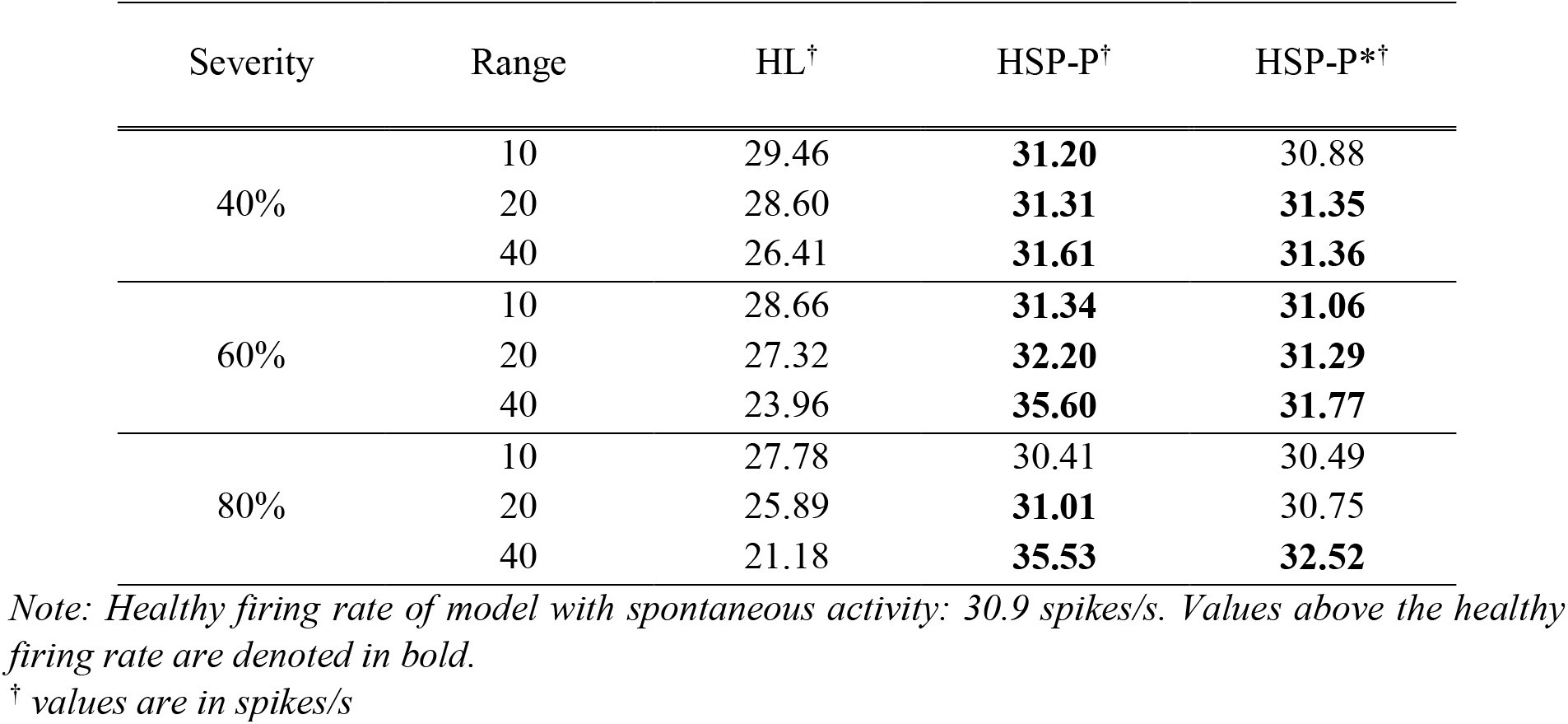
Mean firing rate of models with spontaneous activity in response to a broadband sound.

| Severity | Range | HL<sup>†</sup> | HSP-P<sup>†</sup> | HSP-P*<sup>†</sup> |
| --- | --- | --- | --- | --- |
| 40% | 10 | 29.46 | **31.20** | 30.88 |
|  | 20 | 28.60 | **31.31** | **31.35** |
|  | 40 | 26.41 | **31.61** | **31.36** |
| 60% | 10 | 28.66 | **31.34** | **31.06** |
|  | 20 | 27.32 | **32.20** | **31.29** |
|  | 40 | 23.96 | **35.60** | **31.77** |
| 80% | 10 | 27.78 | 30.41 | 30.49 |
|  | 20 | 25.89 | **31.01** | 30.75 |
|  | 40 | 21.18 | **35.53** | **32.52** |

*Note:* Healthy firing rate of model with spontaneous activity: 30.9 spikes/s. Values above the healthy firing rate are denoted in bold.
<sup>†</sup> values are in spikes/s

**Figure 6.**
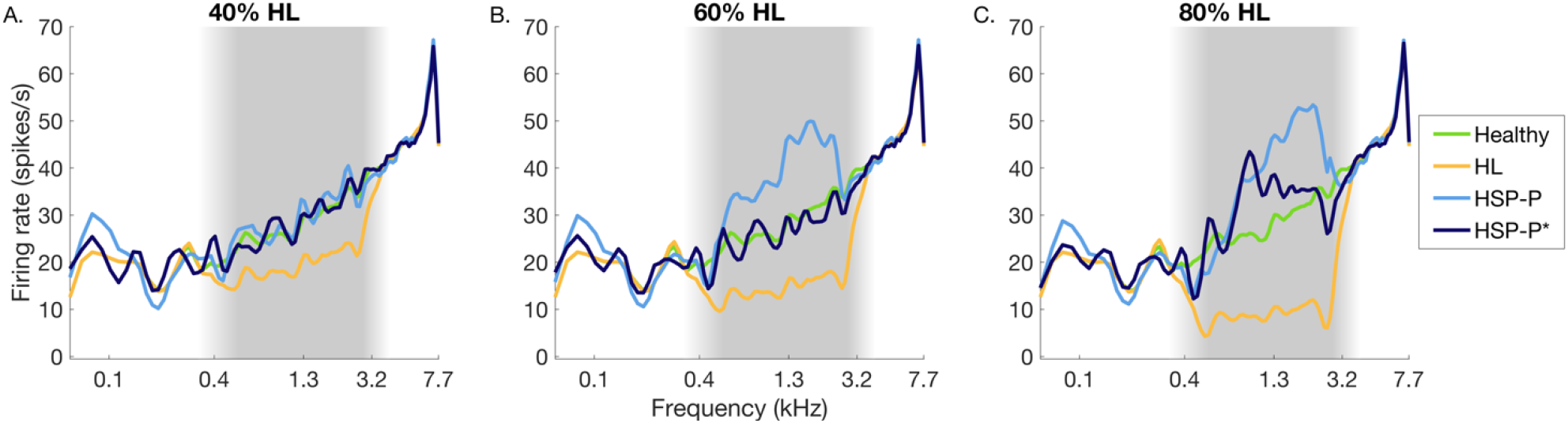
Model responses with spontaneous activity to broadband noise. Sound-evoked response in the healthy, HL, and HSP without (HSP-P) and with updated thalamocortical input (HSP-P*) are shown in green, yellow, light and dark blue, respectively. A, B and C show the responses for models with 40%, 60% and 80% HL, respectively. For all models, HL affected 40 units. Shaded grey regions delineate the HL region.

### 3.4 HSP Broadens Frequency Tuning and Increases Neural Noise

To explore how HSP affected frequency preference and selectivity, FTCs were constructed by recording each unit’s response to a range of pure tones in models with spontaneous activity. *Figure 7* shows the average FTCs outside, inside the lower sloping region, and within the HL region. Note that in the healthy model, in accordance with results from animal electrophysiology (Cheung et al., 2001; Imaizumi et al., 2004) and the existing model implementation (Zulfiqar et al., 2020) an increase in frequency selectivity (i.e., a decrease in tuning width [TW]) with increasing frequency preference (best frequency [BF]) of a unit was present. As TW decreased exponentially with increasing BF, TW in the healthy model was broadest outside the hearing loss region, slightly narrower in the lower sloping HL region, and most narrow inside the HL region.

**Figure 7.**
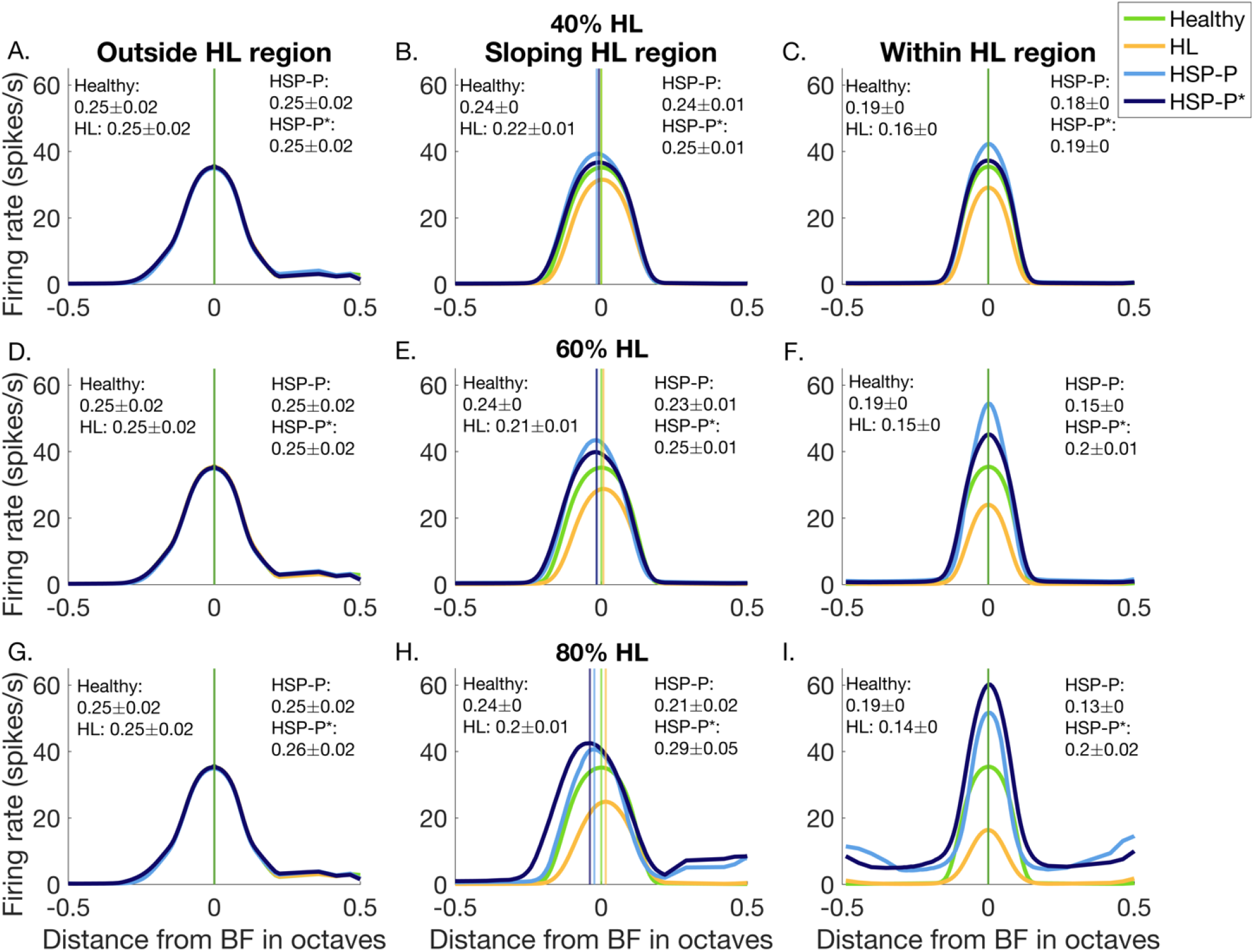
HSP-induced changes in frequency preference and selectivity in models with spontaneous activity. Average FTCs for units outside, in the sloping and within the HL region. Mean TW (in octaves) ±SEM is shown for the different models. Vertical lines indicate the average best frequency (BF) for the different models. A, D, and G show average FTCs of units outside the HL region for 40%, 60% and 80% HL models, respectively. B, E, and H show average FTCs of units within the sloping HL region for 40%, 60% and 80% HL models, respectively. C, F, and I show average FTCs of units within the HL region for 40%, 60%, and 80% HL models, respectively. Green: healthy, yellow: HL, light blue: FTC of models where P was not updated (HSP-P), dark blue: FTC of the model where P was updated (HSP-P*).

Overall, neither HL nor HSP affected FTC shape outside the HL region (left column of *Fig. 7*). HL reduced the sound-driven response of units in the sloping HL and HL regions, as indicated by the lower firing rate to the unit’s BF as well as neighboring frequencies, resulting in TW narrowing (increased frequency selectivity after HL). Inside the sloping HL and within the HL region, both HSP implementations restored firing rate and even caused an overcompensation in the response to pure tones compared to the healthy model. As expected, the increased excitatory drive and decreased inhibition after HSP resulted in a broader TW (i.e., reduced frequency selectivity; *Fig. 7*) compared to the HL scenario. This change was especially prominent for higher HL scenarios and for the HSP-P* case, where TW became even broader than that in the healthy model. Interestingly, in the sloping and within the HL region of the 80% HL model, we furthermore observed an increased responsiveness to non-preferred frequencies. This pattern was present for both HSP implementations, and may reflect increased neural noise. Comparable observations could be made for the other HL ranges (see. *Fig. S8 and S9*).

Additionally, we observed a minor shift in the unit’s frequency preference (i.e., best frequency [BF]) that was limited to the sloping HL region (see vertical lines in *Fig. 7B, 5E, 5H)*. HL resulted in a shift of the BF to higher frequencies, whereas both HSP mechanisms reduced the BF compared to the healthy model. While the shift for models with updated cortico-cortical connectivity (HSP-P) was similar across all HL severities, models with additionally updated thalamocortical input (HSP-P*) showed a greater shift for more severe HL scenarios.

Finally, in order to explore the origin of the HSP-induced changes in frequency preference and selectivity, we also constructed FTCs in the model without spontaneous activity. We observed only two differences compared to the model with spontaneous activity. Comparable observations were made across HL ranges (*Fig. S10* and *S11*). First, TW in the model without spontaneous activity was overall slightly narrower (i.e., lower TW; compare Fig. 7 to Fig. 8) and HSP did not fully compensate for the HL-induced narrowing of tuning width. Second, we no longer observed the increased responsiveness to non-preferred frequencies for the case of 80% HL. Thus, the presence of spontaneous activity in the model seems to be a prerequisite for observing increased neural noise after HSP.

**Figure 8.**
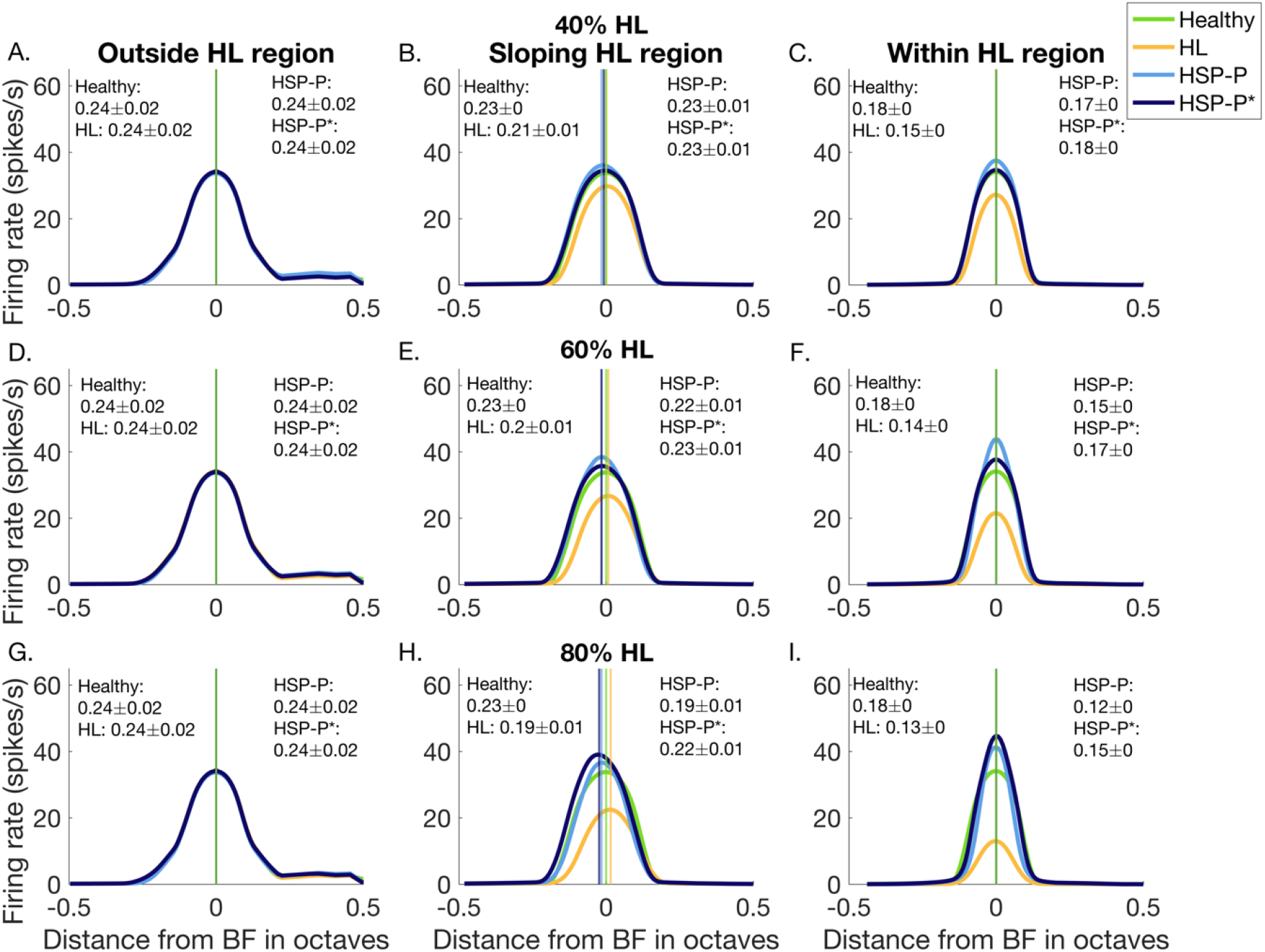
HSP-induced changes in frequency preference and selectivity in models without spontaneous activity. Average FTCs for units outside, in the sloping and within the HL region. Mean TW (in octaves) ±SEM is shown for the different models. Vertical lines indicate the average best frequency (BF) for the different models. A, D, and G show average FTCs of units outside the HL region for 40%, 60% and 80% HL models, respectively. B, E, and H show average FTCs of units within the sloping HL region for 40%, 60% and 80% HL models, respectively. C, F, and I show average FTCs of units within the HL region for 40%, 60%, and 80% HL models, respectively. Green: healthy, yellow: HL, light blue: FTC of models where P was not updated (HSP-P), dark blue: FTC of the model where P was updated (HSP-P*).

## 4. Discussion

We implemented HSP in a firing rate model of the auditory cortex and explored the influence of homeostatic principles after peripheral hearing loss, hypothesized as the causal mechanism underlying tinnitus, on model responses. We observed that HSP restored the model’s firing rate but also resulted in neural signatures previously associated with tinnitus. That is, we observed elevated spontaneous and sound-driven activity, spatiotemporal oscillations, and enhanced neural noise. Below we interpret these observations in light of experimental findings across species and brain imaging techniques.

Increased spontaneous activity in the auditory cortex has been repeatedly observed in animal models of tinnitus (Seki and Eggermont, 2003) and previous computational models of HSP (Chrostowski et al., 2011; Schaette and Kempter, 2006). In our model, the increased spontaneous activity organized in spatiotemporal oscillations. The WCCM represents a model of two diffusively coupled excitable systems (i.e., the excitatory and inhibitory subpopulation). As such, the model can exhibit multiple different spatiotemporal dynamics, or modes, depending on the parameter selection (Bressloff, 2012). In the model of healthy auditory processing, parameters were chosen such that recurrent model excitation activated the inhibitory response reducing the overall activity of the model (i.e., active transient mode; Zulfiqar et al. (2020)). Under HSP conditions, however, model excitability in the HL region was increased. Increased excitability combined with low activity levels is known to result in spatially periodic activity patterns known as Turing instability (Ermentrout and Cowan, 1979; Negahbani et al., 2015). In accordance with previous reports, in our model output Turing oscillations were limited to the HL region with increased excitability and were observed only in low-activity states. That is, in the presence of the broadband noise, the increased input prevented model instability. Although Turing instability has previously been linked to visual hallucinations (Ermentrout and Cowan, 1979), there is no evidence for a spatially organized activity pattern in the auditory cortex of tinnitus patients. However, the existence of spatial organization in spontaneous activity is difficult to assess through human neuroimaging, as the spatial resolution of magnetoencephalography (MEG) and electroencephalography (EEG) is insufficient, and the scanner noise during data acquisition prohibits observing the auditory system at rest using fMRI. It may be of interest to explore changes in metabolic demand with tinnitus (e.g., cerebral blood flow or cerebral metabolic rate of glucose), reflecting spontaneous neural activity, through arterial spin labeling (ASL) or positron emission tomography (PET; Chen et al. (2015)). At a behavioral level, the loudness of the tinnitus percept can be reduced by background noise or masked by auditory input (Fournier et al., 2018). The disappearance of Turing instability with auditory drive could represent this finding. In follow up work it would be of interest to detail the extent to which the appearance of Turing oscillations in model output matches psychoacoustic findings in tinnitus patients.

The observed temporal oscillations in model responses after HSP, varying from 27 Hz to 45 Hz, likely resulted from a Hopf instability (Negahbani et al., 2015). MEG and EEG studies have shown that tinnitus is accompanied by increases in gamma band activity (> 30 Hz) within the auditory cortex (van der Loo et al., 2009). Further analysis of the model’s temporal response during spontaneous activity is needed to relate the presence of Hopf oscillations in the model to tinnitus. Specifically, it may be of interest to directly test the model prediction that the oscillation frequency depends on HL severity. If evidence of such dependence is observed, fine-tuning model oscillation rate to the frequency observed in individual patients could suggest inter-individual differences in tinnitus neurobiology.

The increased sound-driven response after HSP is in line with empirical findings made from the auditory cortex in animal models of tinnitus (Kimura and Eggermont, 1999; Seki and Eggermont, 2003). Sound-driven activity with tinnitus has been extensively explored with fMRI, and resulted in mixed findings. Hyperactivity in the auditory cortex of tinnitus patients was reported by a number of studies (Gu et al., 2010; Leaver et al., 2011), but these findings did not hold when correcting for hearing loss and hyperacusis (Koops et al., 2020). Studies with control measures for these confounding comorbidities reported no difference in the auditory cortical sound-driven response with tinnitus (Berlot et al., 2020; Koops et al., 2020) or even observed a decreased response in tinnitus patients compared to controls (Hofmeier et al., 2018; van Gendt et al., 2012). Thus, while increased sound-driven activity within the auditory cortex was traditionally seen as hallmark of tinnitus, it has more recently been proposed to instead reflect co-occurring hyperacusis (Knipper et al., 2013). We implemented HSP to restore the firing rate to broadband noise in a model without spontaneous activity. This led to an overcompensation in response to pure tones and to all sounds in a model with spontaneous activity. This is an interesting finding in itself, as it shows that HSP-driven response restoration, and whether or not overcompensation results from it, depends on the stimulus that HSP is based on. Taking into account the fMRI literature, it may also suggest that our HSP implementation (i.e., based on responses to broadband noise) represents hyperacusis rather than tinnitus. Instead, HSP based on responses to narrowband sounds may better reflect the neurobiology underlying tinnitus, as it would lead to smaller changes in the synaptic weights, thus preventing overcompensation.

As our model covered a full tonotopic axis, we could evaluate HSP-induced changes in frequency preference and selectivity. Through increased excitation and decreased inhibition, HSP resulted in a broader tuning width (i.e., decreased frequency selectivity) compensating for the TW narrowing that resulted from HL. Overcompensation was especially apparent for higher intensities and ranges of hearing loss, where the TW after HSP became broader than that of the healthy model. This result is in accordance with a previous auditory cortical model of HSP (Chrostowski et al., 2011) and electrophysiological studies (Schreiner et al., 2000). TW changes have not been observed in fMRI studies of tinnitus. However, the spectral resolution with which fMRI studies have assessed frequency selectivity was too coarse to pick up the HSP-induced TW changes predicted by our model (i.e., below 0.05 octaves). Our results suggest that it would be of interest to explore TW changes in tinnitus patients with high spectral resolution. Through natural sound stimulation combined with PRF mapping or encoding (Dumoulin and Wandell, 2008; Santoro et al., 2014; Thomas et al., 2015), a high spectral resolution of frequency selectivity measurements could be achieved in a time-efficient manner.

We did not observe major changes in the frequency preference of the model units, suggesting that HSP itself is insufficient for eliciting large-scale tonotopic reorganization. The lack of tonotopic reorganization with tinnitus is in accordance with recent fMRI studies in tinnitus patients (Berlot et al., 2020; Koops et al., 2020; Langers et al., 2012), whose findings suggested that earlier results of tonotopic map changes (Mühlnickel et al., 1998; Rauschecker, 1999), resulted from HL rather than underlying tinnitus. Only in the lower sloping region, small BF changes (< 0.1 octaves) were observed. That is, while HL resulted in a shift of the best frequency to higher frequencies (i.e., towards the HL region), the best frequency of units after HSP shifted downwards (i.e., away from the HL region). The direction of BF shift can be explained by the difference in response strength of units above vs. below the lower sloping region. After HL, the response to pure tones was stronger in units below the sloping region (as hearing loss reduced the response in units above the sloping region). Units in the sloping region therefore received stronger lateral inhibition from lower compared to higher units, increasing the BF of units within the sloping region. Instead, after HSP the response to pure tones in units above the sloping region was strongest, resulting in stronger lateral inhibition from units above the sloping region and shifting the best frequency of units within the sloping HL region to lower frequencies. While our model findings suggest that it would be of interest to map frequency preference at the HL edge of tinnitus patients with fMRI to test for BF changes, this neural signature is unlikely to be the source of tinnitus. That is, if these predicted BF changes would underlie tinnitus, the tinnitus pitch would be expected to be within the normal-hearing frequency range or near the frequency edge. Instead, the tinnitus pitch is generally located within the HL region (Schaette and Kempter, 2009; Yang et al., 2011).

Finally, for high levels of hearing loss, HSP resulted in increased model responses to non-preferred frequencies. Interestingly, this observation is in line with recent ultra-high field fMRI findings where stronger responses to non-preferred frequencies were seen in the inferior colliculus and auditory cortex of tinnitus patients compared to hearing loss-matched controls. This finding was interpreted as evidence of reduced inhibition and increased neural noise (Berlot et al., 2020). In accordance, we only observed increased responses to non-preferred frequencies when the model contained spontaneous activity, suggesting that this finding results from an HSP-induced amplification of spontaneous model activity.

Thalamocortical input, the feedforward signal from the medial geniculate body (MGB) to the primary auditory cortex, has been suggested as source of the tinnitus percept. For example, the noise cancellation hypothesis proposes that in the healthy system the MGB filters out tinnitus noise on its way to the auditory cortex. Through MGB disinhibition this filtering function is proposed to fail, resulting in tinnitus (Rauschecker et al., 2010). The implementation of two different HSP mechanisms, either exclusively updating the cortico-cortical synaptic weights (HSP-P) or additionally updating thalamocortical input (HSP-P*) allowed differentiating the relative contribution of cortical and thalamocortical mechanisms to HSP-induced neural signatures. Model behavior was similar across the two HSP implementations, as sound driven responses were restored through both implementations, but minor differences could be observed. Overall, the additional increased input from the thalamus in the HSP-P* implementation brought the model closer to the behavior of the healthy model. That is, as synaptic weights were not updated through amplification of thalamic input, this modification did not intrinsically alter model responses. However, after adding spontaneous activity to the model, HSP with updated thalamocortical input (HSP-P*) resulted in stronger tinnitus-related signatures than when only cortico-cortical synaptic weights were updated (HSP-P). This was especially observed for models with high HL range and intensity, and suggests that increasing thalamocortical input can enhance neural noise through amplification of spontaneous activity. Our model results thereby put forward a neurobiological mechanism through which the noise cancellation hypothesis (Rauschecker et al., 2010) could operate.

In addition to experimentally validating the model predictions outlined above, it would be of interest to extend the proposed model in future work. For example, while we implemented homeostatic mechanisms after HL by altering the strength of synaptic connections, other mechanisms of neural activity restoration through HSP have been proposed. For instance, increased excitability (i.e., a depolarized resting membrane potential) of auditory cortical pyramidal neurons has been observed after HL (Kotak et al., 2005). By extending the current model implementation to a multiscale framework, where changes in membrane potential at a smaller spatial scale could affect model parameters at the WCCM level, such alternative HSP mechanism could be implemented and tested. Furthermore, it may be of interest to extend the model with a biologically inspired ear module such that interindividual differences in peripheral hearing loss may be modeled (Gault et al., 2018; Zilany et al., 2014), or embed computational models that focus on the tinnitus percept (Gault et al., 2020; Hu et al., 2021). Finally, within the dynamic causal modeling (DCM) framework (Friston et al., 2003), nonlinear biophysical generative models have been proposed that capture the translation from neuronal responses to non-invasive neuroimaging data (a hemodynamic model for fMRI: Havlicek et al. (2015); Stephan et al. (2008), and a lead field model for EEG and MEG: Kiebel et al. (2006)). Extending the current model with such physiological models would allow predicting neuroimaging data through forward modeling (Zulfiqar et al., 2021). Conversely, model inversion would allow making inferences about neural responses across spatial scales based on group differences between tinnitus patients and controls in non-invasive imaging data. In summary, the proposed firing rate model of HSP could serve as a framework to further investigate, through model extension towards smaller and larger spatial scales, the cortical mechanisms underlying tinnitus by means of computational modeling.

## Acknowledgements

This work was supported by the Netherlands Organization for Scientific Research (NWO; VENI grant 451-15-012 to M.M.)

## Code availability statement

All code and output is available at: https://github.com/macsbio/model-homeostaticplasticity-tinnitus

