## Supplementary Material for "Modelling homeostatic plasticity in the auditory cortex results in neural signatures of tinnitus"

This file includes:

- Supplementary Equations S1-S3
- Supplementary Table S1
- Supplementary Figures S1-S11

### Supplementary Equations S1-S3

For updating $w_{EE}$:

$$\tau_{HSP}\frac{dw_{EE_{mn}}}{dt} = \left(r_{target} - r_i(t)\right) * w_{EE_{mn}} \qquad \text{(eq. S1)}$$

For updating *P:*

$$\tau_{HSP}\frac{dw_{a_{m,n}}}{dt} = \left(r_{target} - r_i(t)\right) * w_{a_{mn}} \qquad \text{(eq. S2)}$$

For updating $w_{IE}$:

$$\tau_{HSP}\frac{dw_{IE_{mn}}}{dt} = -\left(r_{target} - r_i(t)\right) * w_{IE_{mn}} \qquad \text{(eq. S3)}$$

**Supplementary Table S1**. Model parameters by Zulfiqar et al. (2020)

| Parameters | Values |
|---|---|
| $M$ | 100 |
| $\theta$ excitation | 60 |
| $\theta$ inhibition | 80 |
| $b_{EE}$ | 1.5 |
| $b_{IE} = b_{EI}$ | 1.3 |
| $b_{II}$ | 1.5 |
| $\sigma_{EE}$ | 40 |
| $\sigma_{IE} = \sigma_{EI}$ | 160 |
| $\sigma_{II}$ | 10 |
| $\tau$ (ms) | 10 |

$M$ is the maximum spike rate, $\theta$ is the semi-saturation constant. Parameters $b_{EE}$, $b_{II}$, $b_{EI}$ and $b_{IE}$ represent the maximum synaptic strength between excitatory units, between inhibitory units, from excitatory to inhibitory units, and vice versa, respectively. Time constant $\tau$ and spatial spread parameter $\sigma$ (EE, II, EI/IE) are listed.

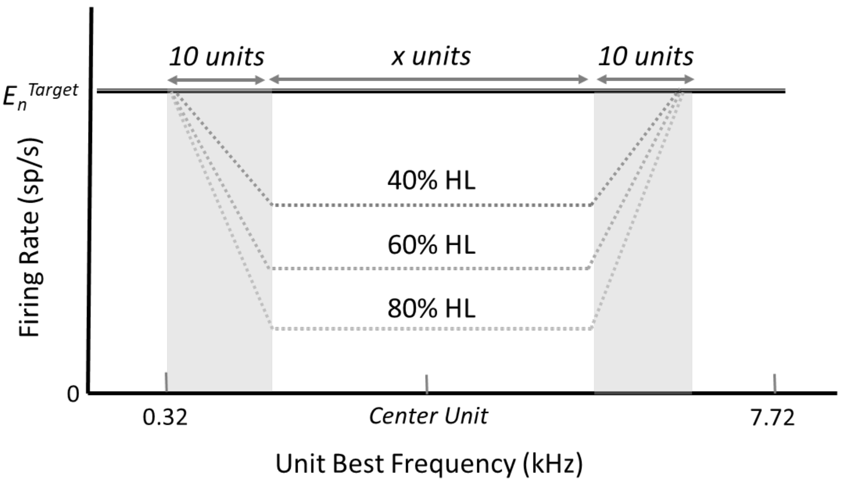


**Supplementary Figure S1. Hearing loss implementation.** Hearing loss was implemented by decreasing the model's filterbank response (firing rate in spikes/s) to a Gaussian white noise signal. $E_n^{Target}$ refers to the response of the healthy model. Across the modeled HL ranges and severities, HL started at 0.32 kHz corresponding to the best frequency (BF) of unit 20. HL severity linearly increased, and firing rate decreased, over the next 10 units (BF = 0.55 kHz). The firing rate decrease was maintained for a HL range of $x$ = 10, 20 or 40 units, corresponding to a BF of 0.87, 1.31 and 2.82 kHz, respectively. To avoid boundary effects and maintain model stability, a gradual decrease of HL severity (and increase of firing rate back to healthy model output) was implemented over the next 10 units (unit 50 [BF = 1.31 kHz], unit 60 [BF = 1.94 kHz] or unit 80 [BF = 4.07 kHz], for HL range 10, 20 and 40, respectively).

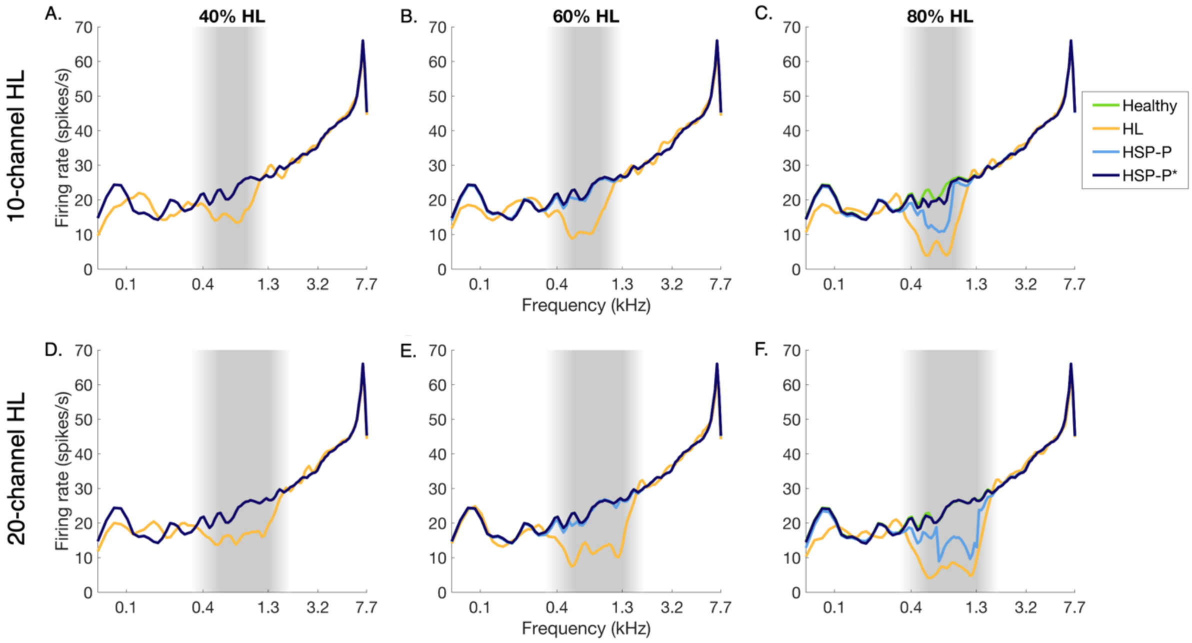


**Supplementary Figure S2. Model responses to broadband noise.** Sound-evoked response in the healthy, HL, and HSP without (HSP-P) and with (HSP-P*) updated input are shown in green, yellow, light and dark blue. Left, middle, and right columns show the responses for models with 40%, 60% and 80% HL, respectively. HL affecting 10 and 20 units are shown in the top and bottom row, respectively. Shaded grey regions delineate the HL region.

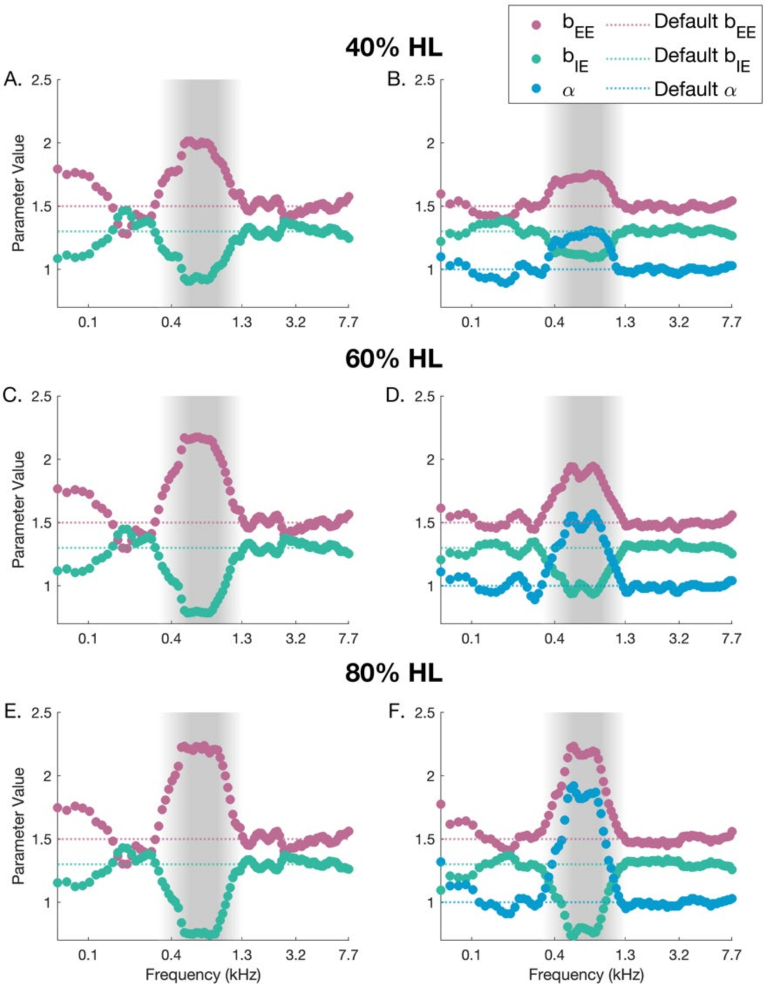


**Supplementary Figure S3. HSP-induced changes in model parameters for 10 channel hearing loss.** Changes in cortico-cortical synaptic weights $b_{EE_{mn}}$ and $b_{IE_{mn}}$ and the multiplication factor $\alpha_n$ for thalamocortical input P are shown across model units. Dark blue: $b_{EE_{mn}}$; turquoise: $b_{IE_{mn}}$; light blue: $\alpha_n$. The dotted lines show the default $b_{EE_{mn}}$, $b_{IE_{mn}}$, and $\alpha_n$ values of the healthy model. A, C, and E show the results for models where thalamocortical input P was not updated (HSP-P), while B, D and F display the values for models where P was updated (HSP-P*). For all models, HL affected 10 units, outlined by the shaded grey region.

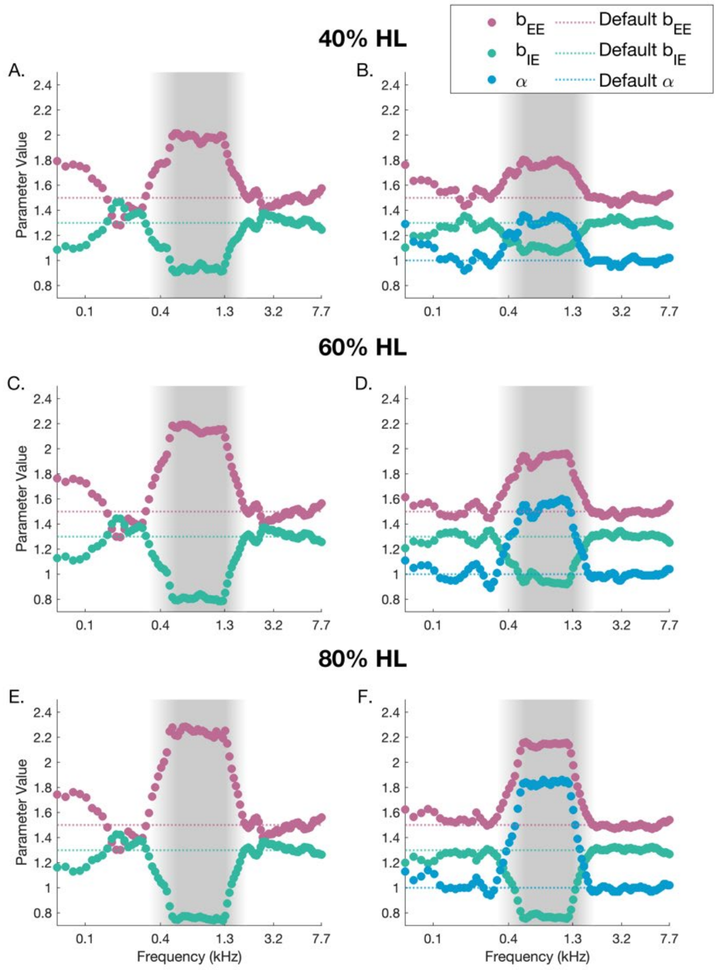


**Supplementary Figure S4. HSP-induced changes in model parameters for 20 channel hearing loss.** Changes in cortico-cortical synaptic weights $b_{EE_{mn}}$ and $b_{IE_{mn}}$ and the multiplication factor $\alpha_n$ for thalamocortical input P are shown across model units. Dark blue: $b_{EE_{mn}}$; turquoise: $b_{IE_{mn}}$; light blue: $\alpha_n$. The dotted lines show the default $b_{EE_{mn}}$, $b_{IE_{mn}}$ and $\alpha_n$ values of the healthy model. A, C, and E show the results for models where thalamocortical input P was not updated (HSP-P), while B, D and F display the values for models where P was updated (HSP-P*). For all models, HL affected 20 units, outlined by the shaded grey region.

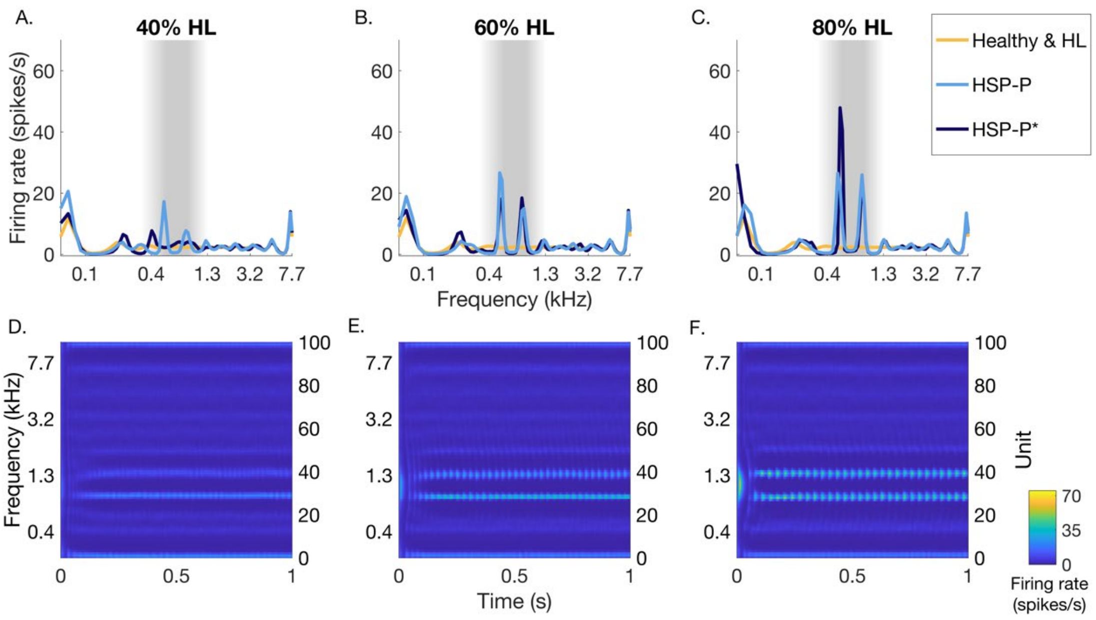


**Supplementary Figure S5. Spontaneous model activity without sound input for 10 channel hearing loss.** Yellow: healthy and HL responses; note that in absence of an input sound these conditions are equal, as spontaneous activity was added after the LIN stage (and was thus not affected by HL) light blue: HSP-P response without updated thalamocortical input; dark blue: HSP-P* response with updated thalamocortical input. A-C show the model responses averaged across time for 40%, 60%, and 80% HL, respectively. D-F show the HSP-P model response over time for 40%, 60%, and 80% HL, respectively. For all models, HL affected 10 units.

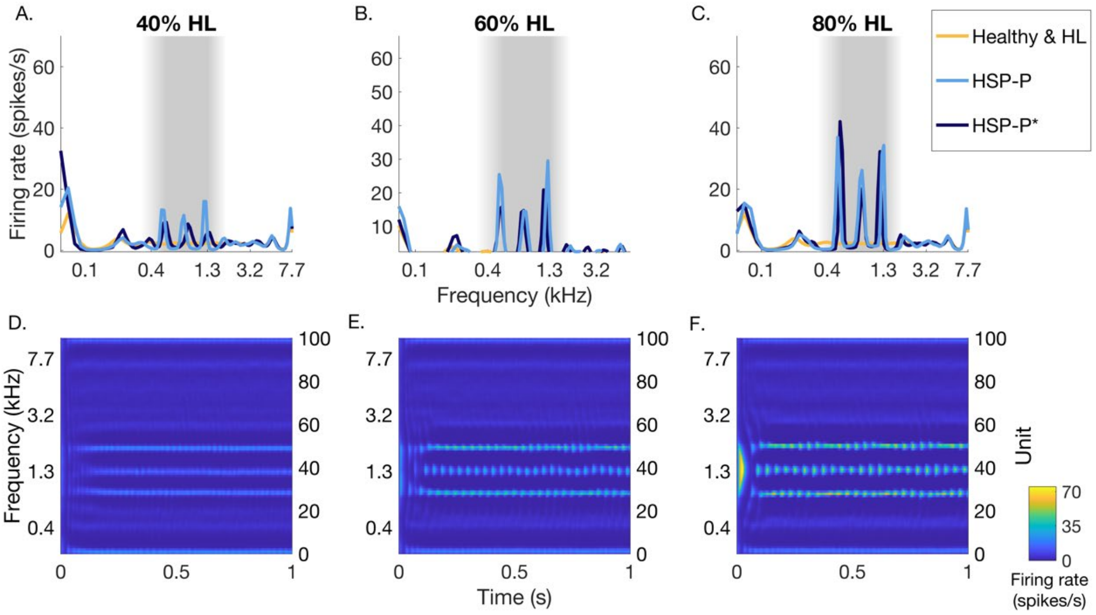


**Supplementary Figure S6. Spontaneous model activity without sound input for 20 channel hearing loss.** Yellow: healthy and HL responses; note that in absence of an input sound these conditions are equal, as spontaneous activity was added after the LIN stage (and was thus not affected by HL) light blue: HSP-P response without updated thalamocortical input; dark blue: HSP-P* response with updated thalamocortical input. A-C show the model responses averaged across time for 40%, 60%, and 80% HL, respectively. D-F show the HSP-P model response over time for 40%, 60%, and 80% HL, respectively. For all models, HL affected 20 units.

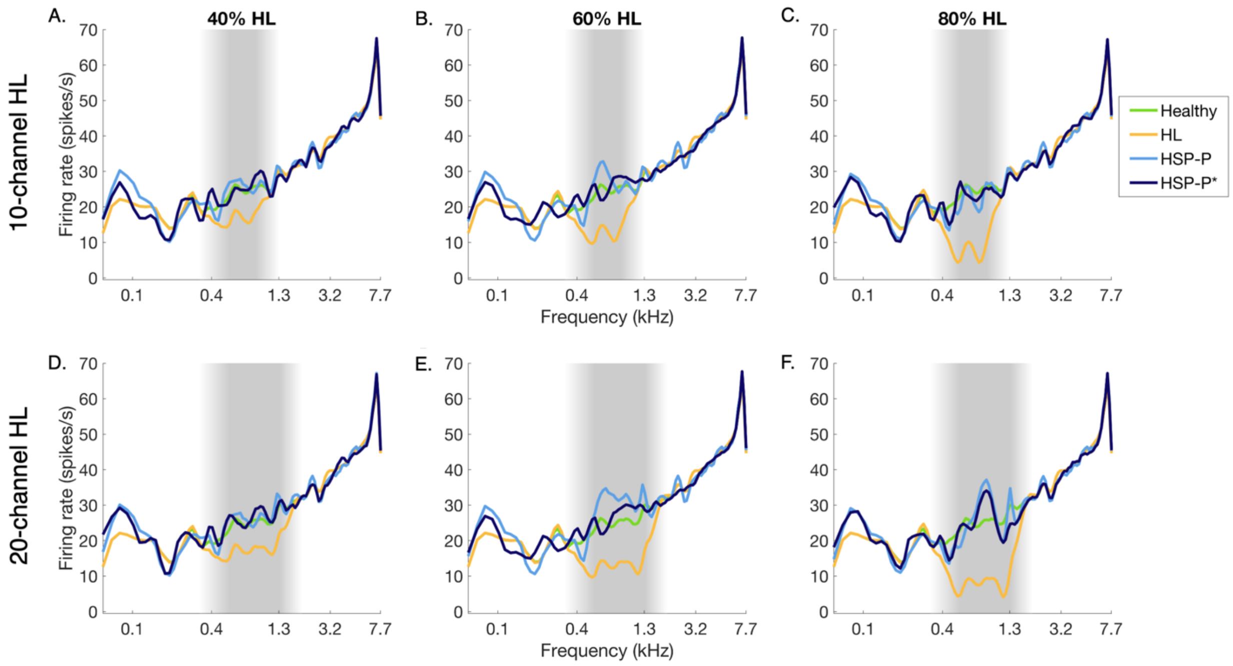


**Supplementary Figure S7. Model responses to broadband noise.** Responses in the healthy model, for HL, HSP without (HSP-P) and with (HSP-P*) updated thalamocortical input are shown in green, yellow, light and dark blue. Left, middle, and right columns show the responses for models with 40%, 60% and 80% HL, respectively. HL affecting 10 units is shown in the top row, and HL affecting 20 units is shown in the bottom row. Shaded grey regions delineate the HL region.

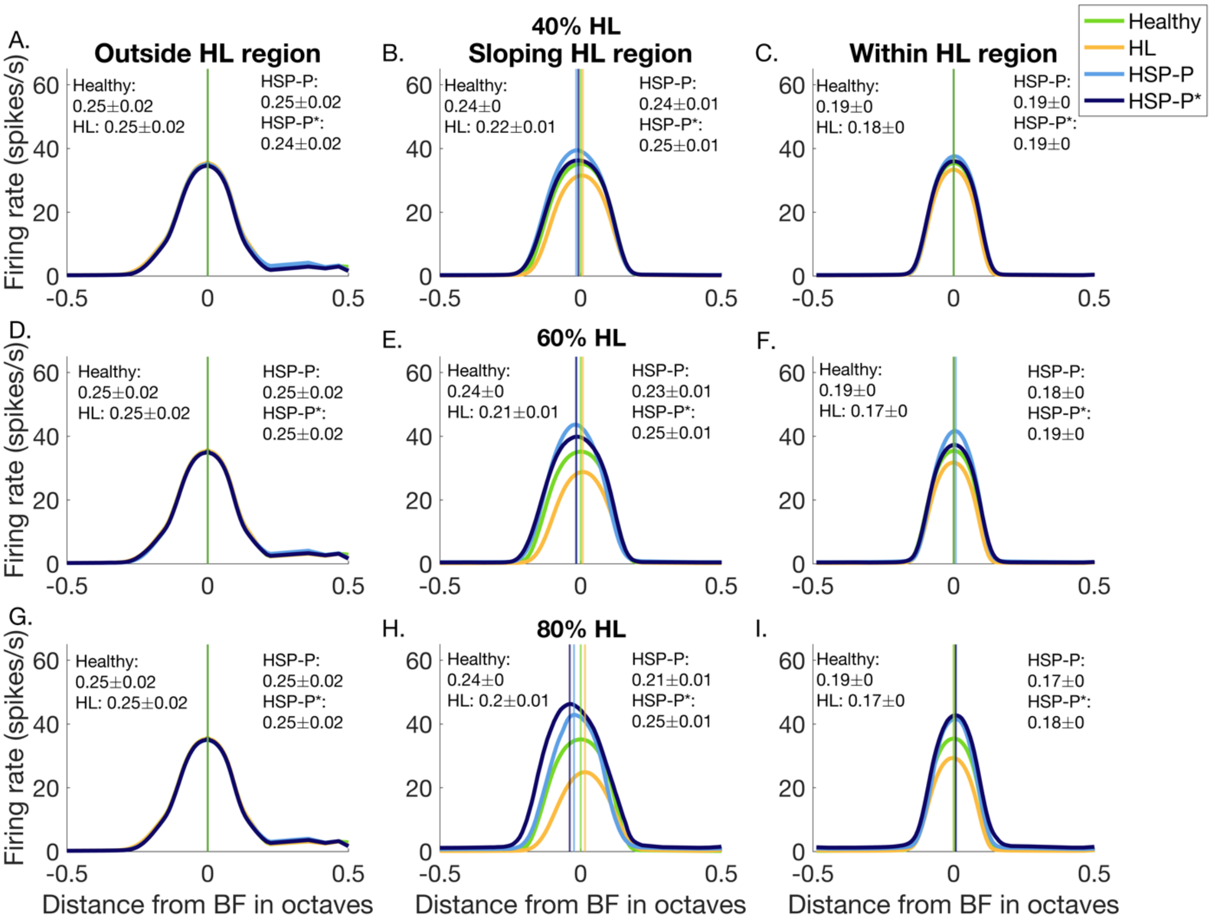


**Supplementary Figure S8. HSP-induced changes in frequency preference and selectivity in models with spontaneous activity for 10 channel hearing loss.** Average FTCs for units outside, in the sloping and within the HL region. Mean TW (in octaves) ±SEM is shown for the different models. Vertical lines indicate the average best frequency (BF) for the different models. A, D, and G show average FTCs of units outside the HL region for 40%, 60% and 80% HL models, respectively. B, E, and H show average FTCs of units within the sloping HL region for 40%, 60% and 80% HL models, respectively. C, F, and I show average FTCs of units within the HL region for 40%, 60%, and 80% HL models, respectively. Green: healthy, yellow: HL, light blue: FTC of models where P was not updated (HSP-P), dark blue: FTC of the model where P was updated (HSP-P*).

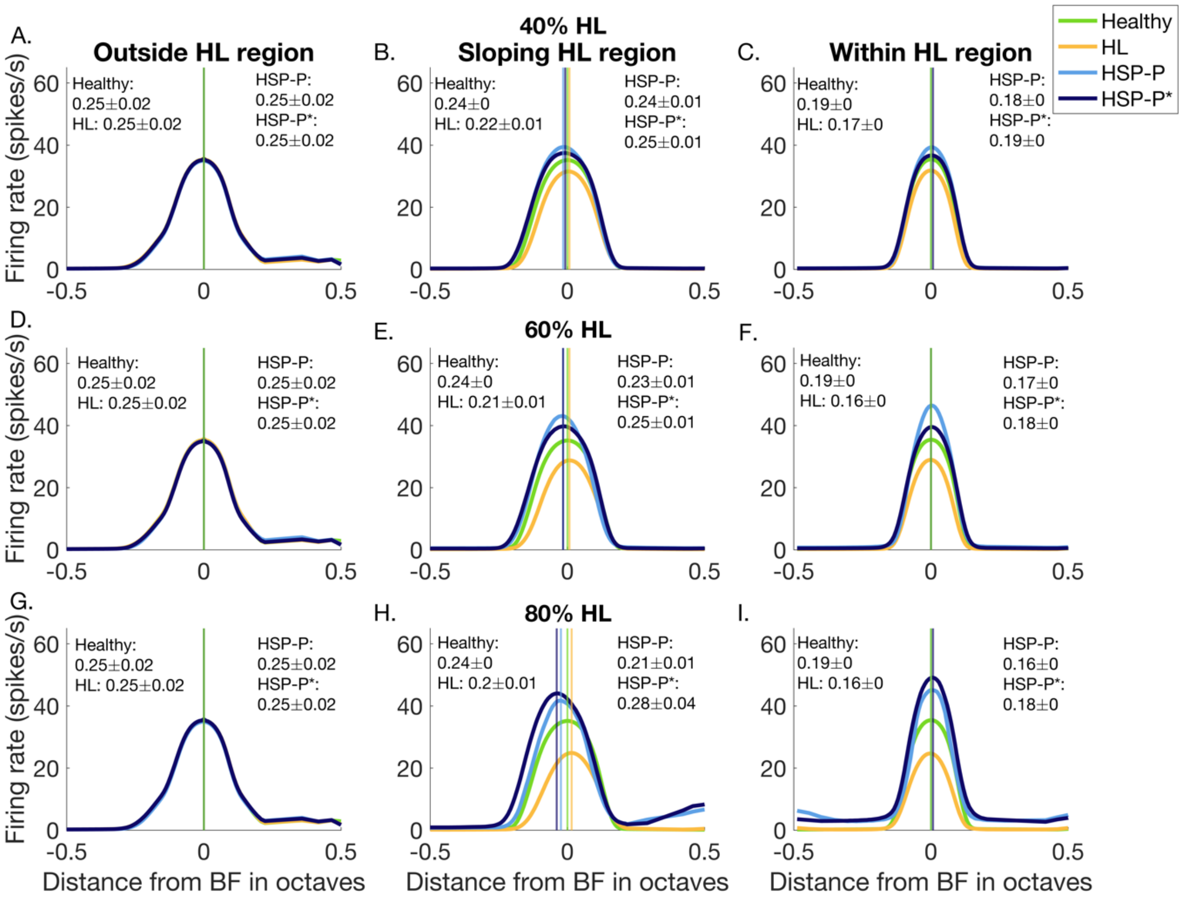


**Supplementary Figure S9. HSP-induced changes in frequency preference and selectivity in models with spontaneous activity for 20 channel hearing loss.** Average FTCs for units outside, in the sloping and within the HL region. Mean TW (in octaves) ±SEM is shown for the different models. Vertical lines indicate the average best frequency (BF) for the different models. A, D, and G show average FTCs of units outside the HL region for 40%, 60% and 80% HL models, respectively. B, E, and H show average FTCs of units within the sloping HL region for 40%, 60% and 80% HL models, respectively. C, F, and I show average FTCs of units within the HL region for 40%, 60%, and 80% HL models, respectively. Green: healthy, yellow: HL, light blue: FTC of models where P was not updated (HSP-P), dark blue: FTC of the model where P was updated (HSP-P*).

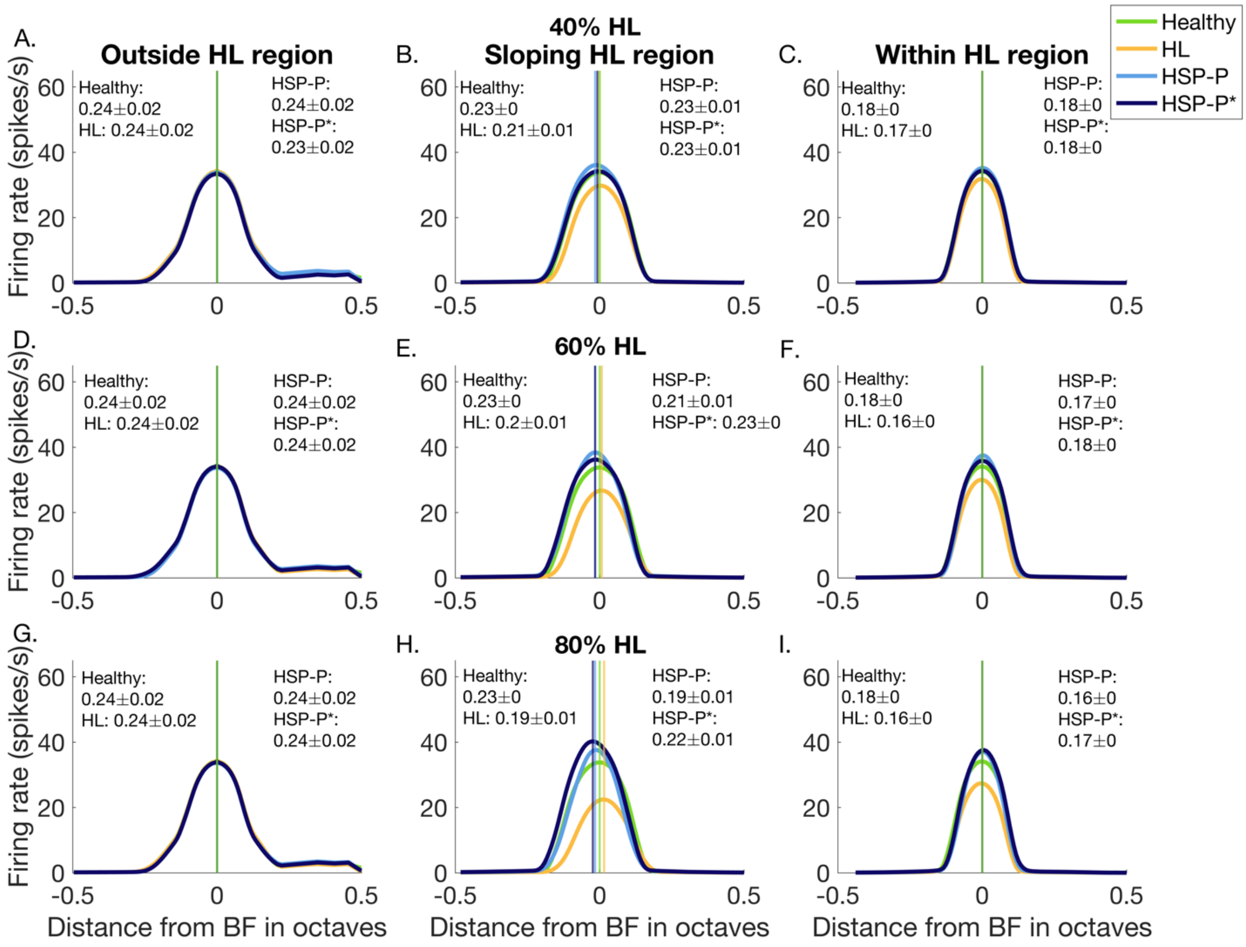


**Supplementary Figure S10. HSP-induced changes in frequency preference and selectivity in models without spontaneous activity for 10 channel hearing loss.** Average FTCs for units outside, in the sloping and within the HL region. Mean TW (in octaves) ±SEM is shown for the different models. Vertical lines indicate the average best frequency (BF) for the different models. A, D, and G show average FTCs of units outside the HL region for 40%, 60% and 80% HL models, respectively. B, E, and H show average FTCs of units within the sloping HL region for 40%, 60% and 80% HL models, respectively. C, F, and I show average FTCs of units within the HL region for 40%, 60%, and 80% HL models, respectively. Green: healthy, yellow: HL, light blue: FTC of models where P was not updated (HSP-P), dark blue: FTC of the model where P was updated (HSP-P*).

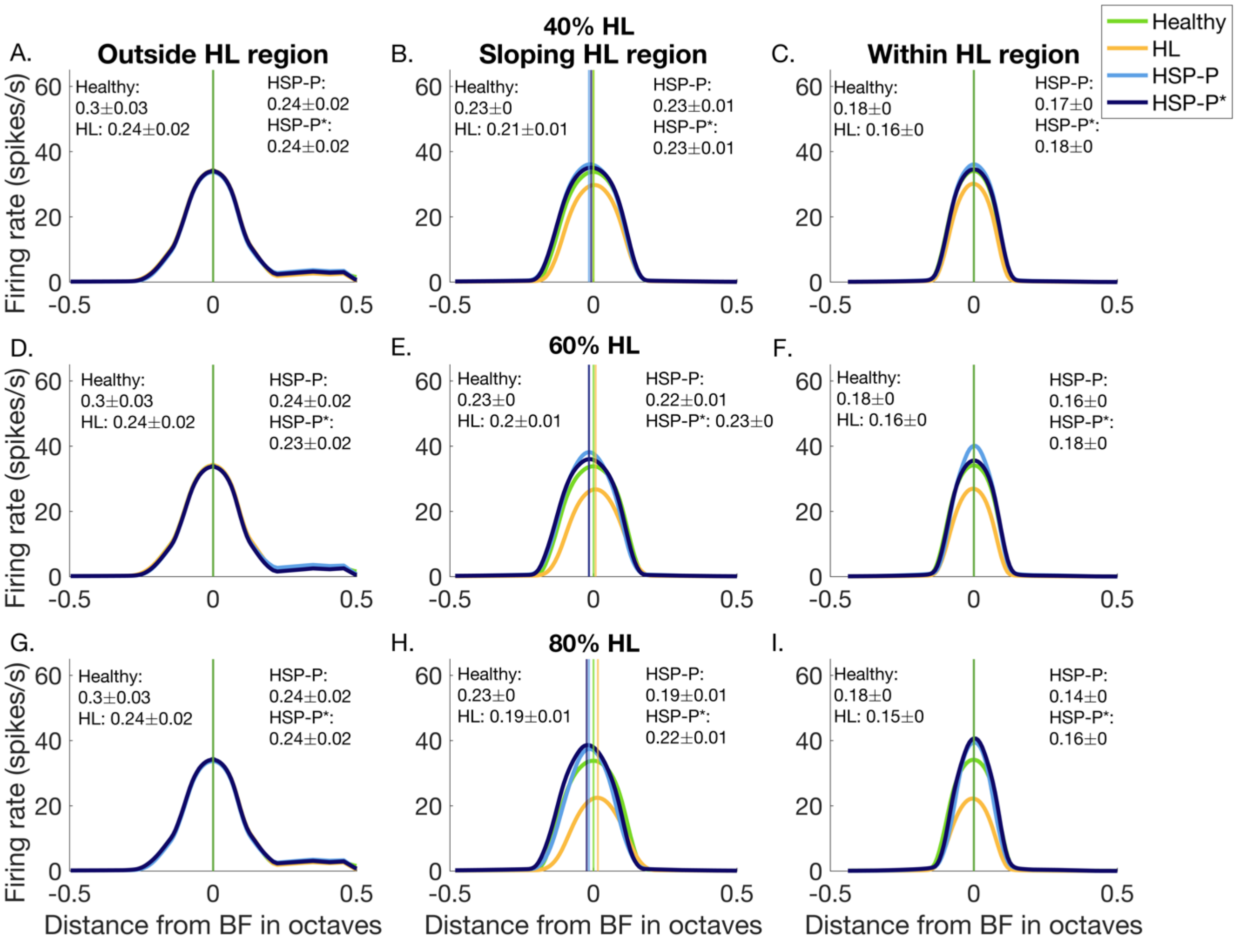


**Supplementary Figure S11. HSP-induced changes in frequency preference and selectivity in models without spontaneous activity for 20 channel hearing loss.** Average FTCs for units outside, in the sloping and within the HL region. Mean TW (in octaves) ±SEM is shown for the different models. Vertical lines indicate the average best frequency (BF) for the different models. A, D, and G show average FTCs of units outside the HL region for 40%, 60% and 80% HL models, respectively. B, E, and H show average FTCs of units within the sloping HL region for 40%, 60% and 80% HL models, respectively. C, F, and I show average FTCs of units within the HL region for 40%, 60%, and 80% HL models, respectively. Green: healthy, yellow: HL, light blue: FTC of models where P was not updated (HSP-P), dark blue: FTC of the model where P was updated (HSP-P*).
